# Dynamic extrinsic pacing of the *HOX* clock in human axial progenitors controls motor neuron subtype specification

**DOI:** 10.1101/2020.06.27.175646

**Authors:** Vincent Mouilleau, Célia Vaslin, Simona Gribaudo, Rémi Robert, Nour Nicolas, Margot Jarrige, Angélique Terray, Léa Lesueur, Mackenzie W. Mathis, Gist Croft, Mathieu Daynac, Virginie Rouiller-Fabre, Hynek Wichterle, Vanessa Ribes, Cécile Martinat, Stéphane Nedelec

## Abstract

Rostro-caudal patterning of vertebrates depends on the temporally progressive activation of *HOX* genes within axial stem cells that fuel axial embryo elongation. Whether *HOX* genes sequential activation, the “*HOX* clock”, is paced by intrinsic chromatin-based timing mechanisms or by temporal changes in extrinsic cues remains unclear. Here, we studied *HOX* clock pacing in human pluripotent stem cells differentiating into spinal cord motor neuron subtypes which are progenies of axial progenitors. We show that the progressive activation of caudal *HOX* genes in axial progenitors is controlled by a dynamic increase in FGF signaling. Blocking FGF pathway stalled induction of *HOX* genes, while precocious increase in FGF alone, or with GDF11 ligand, accelerated the *HOX* clock. Cells differentiated under accelerated *HOX* induction generated appropriate posterior motor neuron subtypes found along the human embryonic spinal cord. The *HOX* clock is thus dynamically paced by exposure parameters to secreted cues. Its manipulation by extrinsic factors alleviates temporal requirements to provide unprecedented synchronized access to human cells of multiple, defined, rostro-caudal identities for basic and translational applications.

## INTRODUCTION

The patterning of bilaterian body is orchestrated by the differential expression of HOX transcription factors along their rostro-caudal axis. cn The initiation of *Hox* gene expression patterns is closely linked to the rostro-caudal extension of the body axis. As axial progenitors contribute to progressively more caudal mesodermal and neuroectodermal structures, the 3’ to 5’ sequence of *HOX* gene activation is translated into a collinear spatial pattern of expression with 3’ *HOX* genes being expressed anteriorly while 5’ *HOX* ones are late-activated and caudally-expressed (Deschamps and Duboule, 2017; Henrique et al., 2015). While extrinsic factors such as Retinoic Acid (RA), Wnt, or Fibroblast Growth Factors (FGFs) play a key role in controlling HOX patterns, cell intrinsic changes in chromatin states within the complexes correlate with the temporal sequence of their transcriptional induction (Bel-Vialar et al., 2002; Deschamps and Duboule, 2017; Liu et al., 2001; Mazzoni et al., 2013; Narendra et al., 2015; Neijts et al., 2017; Philippidou and Dasen, 2013). However, it remains unclear whether the progressive opening of the chromatin along the complexes serves as an internal timer initiated and terminated by extrinsic cues or whether sequences of secreted factors that activate progressively more caudal *HOX* genes define the tempo of the induction (Bel-Vialar et al., 2002; Del Corral and Storey, 2004; Deschamps and Duboule, 2017; Ebisuya and Briscoe, 2018; Lippmann et al., 2015; Mazzoni et al., 2013; Wymeersch et al., 2019). This might result in the current limited control over pluripotent stem cells (PSCs) differentiation into caudal cell types of distinct well-defined rostro-caudal identities (Diaz-Cuadros et al., 2020; Du et al., 2015; Duval et al., 2019; Faustino Martins et al., 2020; Frith et al., 2018; Gouti et al., 2014; Li et al., 2005; Lippmann et al., 2015; Matsuda et al., 2020; Maury et al., 2015; Ogura et al., 2018; Peljto et al., 2010; Verrier et al., 2018). Indeed, the two models imply different strategies to control in vitro PSC differentiation. The “intrinsic model” predicts that the efficient specification of posterior identities will rely on a precise synchronization between the differentiation timing and the internal *HOX* timer. Alternatively, the second mechanism predicts that exposing axial progenitors to the relevant extrinsic cues will entrain the *HOX* clock to generate progenies of defined rostro-caudal identities. To approach this question and its consequences for cell engineering, we thus aimed at generating axial progenitors from human pluripotent stem cells (hPSCs) to study the mechanisms pacing the *HOX* clock during their differentiation into spinal motor neurons (MNs), a cell type that relies on a precise HOX code to acquire appropriate rostro-caudal subtype identities.

## RESULTS

### HOX expression profiles and motor neuron subtypes in human embryonic spinal cord

The differential expression of HOX transcription factors along the vertebrate spinal cord is a major product of *HOX* gene regulation. In spinal motor neurons, this Hox code orchestrates the specification of subtype specific features controlling the formation of locomotor circuits (Dasen, 2017; Philippidou and Dasen, 2013). Whether the spinal HOX code and associated MN subtypes is conserved in human remains unknown preventing faithful assessment of HOX regulation and its link with cell “rostro-caudal” identity during hPSC differentiation. We thus mapped, in human embryos at 6.3 and 7.5 weeks of development, 7 HOX transcription factors that display collinear expression patterns and instruct MN subtype specification in mouse and chick (Philippidou and Dasen, 2013) (Fig. 1A, S1A-D). As in animal models, human MNs expressed ISL1 or HB9 all along the spinal cord (Fig. 1A, circles in Fig. S1A,D) (Amoroso et al., 2013). Within MNs, HOX displayed rostro-caudal patterns resembling those of mouse and chick (Dasen et al., 2003; Liu et al., 2001) : cervical MNs expressed HOXA/C5, while brachial MNs expressed HOXC6, thoracic MNs HOXC9 and lumbar MNs HOXC10. Caudal brachial MNs co-expressed HOXC6 and HOXC8 and anterior thoracic ones HOXC8 and HOXC9. HOXD9 labeled caudal thoracic MNs as well as anterior lumbar MNs together with HOXC10 (Fig. 1A, S1A-D). In amniotes, this Hox code instructs the formation of distinct motor columns that innervate common muscle groups, and motor pools that innervate a single muscle (Philippidou and Dasen, 2013). To be able to assess *in vitro* whether changes in HOX expression result in the specification of appropriate neuronal subtypes, we mapped MN subtype markers in regard to HOX expression domains (Fig 1A, S1A-D). As in mouse and chick, MNs expressing high level of FOXP1 (FOXP1^high^) were observed at brachio-thoracic (HOXC6 and HOXC8/HOXC9) and lumbar (HOXC10) levels (Fig. 1A, S1A-C). They formed a lateral motor column (LMC) where limb innervating MNs are located (Amoroso et al., 2013; Routal and Pal, 1999). Within this FOXP1^high^ LMC, SCIP/HOXC8 MNs were observed in the caudal brachial spinal cord. In contrast, SCIP/HOXC8/HOXC9 MNs were located in the anterior thoracic region. Their location and their transcriptional code identify them as putative hand-controlling MNs (Fig. S1B,C) (Bell et al., 2017; Mendelsohn et al., 2017). Overall, HOX transcription factors are regionally expressed along the rostro-caudal axis of the human spinal cord. Within these different HOX domains, distinct MN subtypes, identifiable by combination of transcription factors, are generated at stereotyped positions. This data provided readouts to assess the mechanisms regulating *HOX* expression in axial progenitors and their impact on cell type specification during hPSC differentiation.

**Figure 1.**
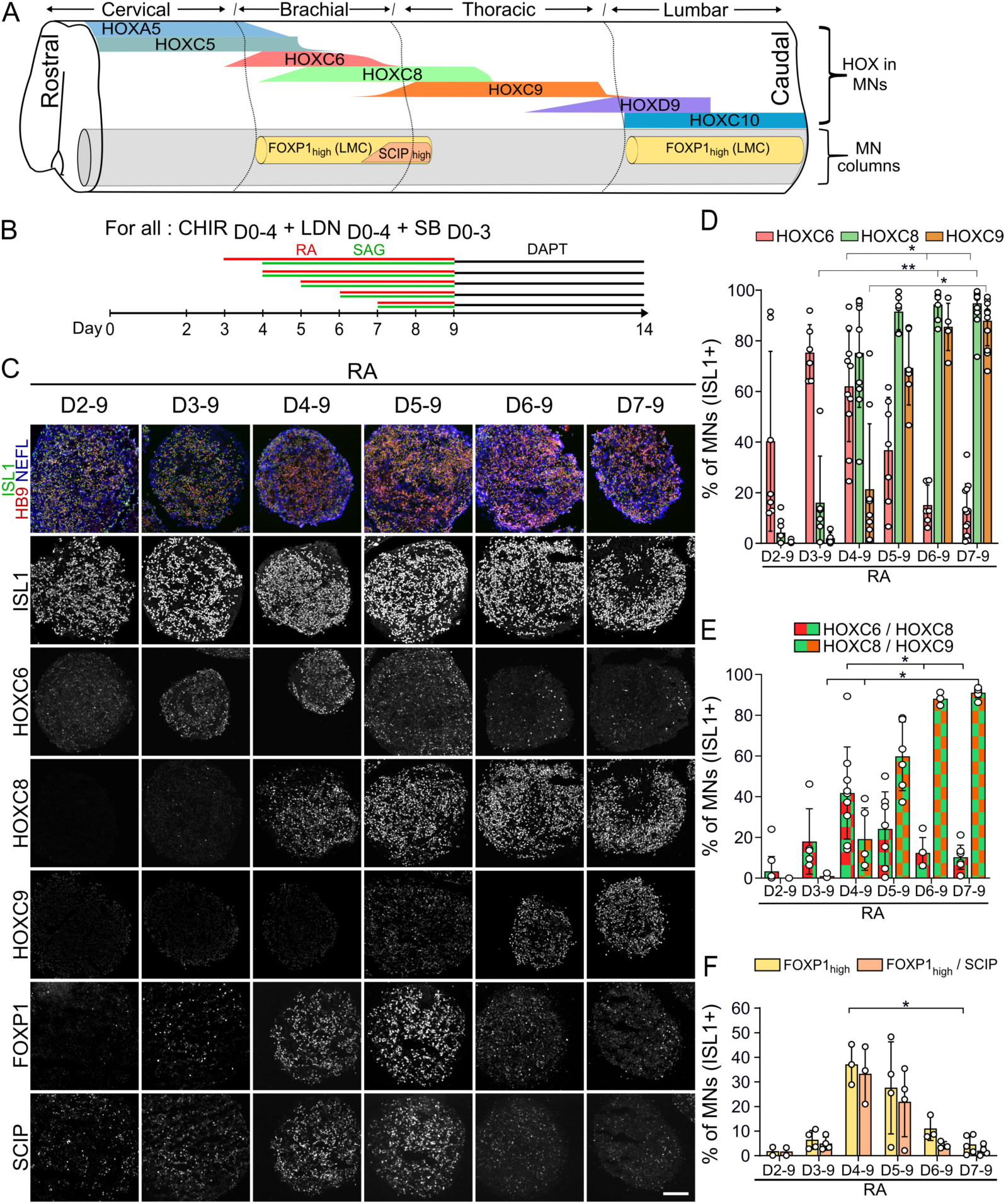
Aging hPSC-derived axial progenitors generate progressively more caudal motor neuron subtypes. **(A)** Schematic summary of data in Fig. S1. MNs defined by the expression of ISL1 or HB9 (here in grey) are organized in motor columns in spinal ventral horns. HOX expression profiles within MNs and localization of MNs expressing high level of FOXP1 and SCIP are represented as observed in the human embryonic spinal cord at 6.3 and 7.5 weeks of gestation. Changes in shapes indicate an increase or decrease in the number of MNs expressing a given marker. FOXP1^high^ MNs are observed selectively in lateral motor columns (LMC) of the brachio-thoracic and lumbar spinal cord. SCIP^high^ MNs are a subset of LMC MNs in the caudal brachial spinal cord. **(B)** Differentiation conditions used in C to F in which the time of exposure to RA/SAG is modulated. **(C)** Immunostaining for ISL1, HB9 (MNs), NEFL (neurons), HOX transcription factors, FOXP1 and SCIP on cryostat sections of EBs on day 14 of differentiation. The later RA is applied the more caudal MNs are. FOXP1 and SCIP MNs are mostly generated in RA D4 and D5 further defining the rostro-caudal identity of the MNs within the HOXC8+ conditions. Scale bar: 100 µm. **(D-F)** Proportion of MNs (ISL1+ cells) expressing the indicated markers. Data are shown as mean ± SD. Each circle is an independent experiment. * if P ≤ 0.05, ** if P ≤ 0.01 and *** if P ≤ 0.001. See also Fig. S1 and S2.

### Aging axial progenitors generate more caudal neuronal subtypes

We first sought to test whether axial progenitors (i.e. progenitors competent to generate caudal cell types of distinct rostro-caudal identities) could be generated using 3D differentiation of human embryonic stem cells (hESC) by monitoring MN subtype specification. We previously reported sequences of extrinsic cues leading to the targeted generation of spinal MNs from human PSCs through a putative axial progenitor stage (Maury et al., 2015). Exposure of embryoid bodies (EBs) to a Wnt pathway agonist (CHIR99021) and inhibitors of BMP and TGFβ pathways (SB431542 and LDN193189) generated progenitors that expressed CDX2, a marker of axial progenitors and a regulator of caudal *HOX* gene induction (Bel-Vialar et al., 2002; Bialecka et al., 2010; Gouti et al., 2014; Maury et al., 2015; Mazzoni et al., 2013; Neijts et al., 2017). Spinal MNs (70% of the cells in average) were generated upon exposure, at day 2 or 4, to Retinoic acid (RA) and an agonist of the sonic hedgehog pathway (SAG) which respectively promotes neurogenesis and guides axial progenitors towards a ventral MN fate (Briscoe and Novitch, 2008; Maury et al., 2015; Ribes et al., 2009). Here, we assessed the rostro-caudal identity of MNs produced in these conditions (Fig. 1B). Staining for HB9, ISL1 and the pan-neuronal marker neurofilament light chain (NEFL) together with quantification of ISL1+ cells, confirmed the efficient generation of spinal MNs as previously shown (Maury et al., 2015) (Fig. 1C, S2A). Analysis of HOX expression showed that RA/SAG from Day 2 (D2) up to D9 gave rise to HOXC6 MNs corresponding to anterior brachial MNs; an identity acquired by most MNs following addition of RA/SAG from D3 to 9 (74.6% of HOXC6, 17.3% HOXC8 MNs) (Fig 1D, E). Addition of RA/SAG from D4 to 9 generated MNs with a caudal brachial identity (41.8% HOXC6/C8) from which 37.1% expressed high level of FOXP1 corresponding to limb-innervating MNs in the spinal cord (Fig.1D-F, S2B). Overall, these results suggested that MN subtype identity was dependent on either i) the duration of exposure to RA; with a shorter RA exposure promoting caudalization or ii) the time at which they received RA as previously suggested (Bel-Vialar et al., 2002; Del Corral and Storey, 2004; Lippmann et al., 2015). To distinguish between these possibilities, D3 progenitors were exposed to reduced duration of RA (D3-5, D3-8 versus D3-9). None of these shorter RA treatments promoted the birth of more caudal MNs. These results showed that the day at which progenitors are exposed to RA/SAG is the main trigger of caudalization (Fig. S2C-E). Further delaying RA/SAG induced even more caudal MN subtypes. On D5, it generated MNs with an anterior thoracic identity (59.8% HOXC8/C9) (Fig. 1B-E) from which 27.6% MNs acquired a FOXP1+ limb-innervating identity (Fig. 1F). HOXC9/FOXP1/SCIP, which are located in the human anterior thoracic spinal cord and might correspond to hand-innervating MNs, were observed almost exclusively in this condition (Fig. 1F, S2F-G). Then RA/SAG (D6 or 7) specified MNs acquiring a mid-thoracic identity as demonstrated by the expression of HOXC9 and the loss of FOXP1^high^ MNs (Fig. 1B-F, S2F-G). The progressive caudalization of MN identity upon incremental delays of RA/SAG addition was confirmed with an induced PSC (iPSC) line. Importantly, an increased concentration of CHIR (from 3 to 4 µM) was necessary to generate a homogenous population of progenitors expressing CDX2 on D3 and to observe the subsequent specification of caudal MN subtypes (Fig. S2H-J). This suggests line to line differences, a result of importance for future studies.

Overall, Wnt activation combined to TGF-β/BMP pathway inhibition converts hPSC into progenitors competent to generate progenies expressing distinct HOX combinations and t found at different rostro-caudal positions in human embryos. Hence, these progenitors qualify as axial progenitors. The duration of the time window between Wnt and RA establishes the final positional identity of the progenies.

### Parallel induction of caudal *HOX* genes and FGF target genes in aging axial progenitors

Then, we sought to determine the molecular changes occurring over time in aging axial progenitors, prior to RA exposure. We performed a comparative transcriptomic analysis of hESC and hESC-derived axial progenitors on day 2, 3 and 4 of differentiation (Fig. 2A and Table S1). Pathway analysis of the genes enriched more than 2 fold (p<0.05) in the progenitors compared to hESC indicated an activation of the Wnt pathway paralleled by a transcriptional activation of *HOX* genes (not shown). In agreement with their axial potential, D2, D3 and D4 progenitors showed a high enrichment in transcripts characterizing the cells of the mouse caudal lateral epiblast in which axial progenitors reside (Expression of *CDX1* and *2, TBXT (BRACHYURY), FGF17, RXRG, SP5/8, WNT5A/B, WNT8A, FGF8, GRSF1, CYSTM1, HES3* together with *SOX2)*; a loss of most pluripotency markers and the absence of typical markers of node cells, mesodermal and allantois (Fig. 2B-C and Table S1) (Cambray and Wilson, 2007; Edri et al., 2019; Gouti et al., 2014; Gouti et al., 2017; Henrique et al., 2015; Koch et al., 2017; Wymeersch et al., 2016; Wymeersch et al., 2019). Furthermore, their transcriptome is highly similar to the one of mouse bipotent neuromesodermal progenitors (NMPs), which represent a population of axial progenitors (Gouti et al., 2014; Gouti et al., 2017; Henrique et al., 2015; Tzouanacou et al., 2009; Wymeersch et al., 2016). Among the 142 genes defining early and late NMPs (Gouti et al., 2017), the D2, D3 or D4 progenitors expressed 122 of them which included the most enriched genes when comparing progenitors to hESC (Fig. 2C). Accordingly, D2 and D3 axial progenitors co-expressed SOX2, CDX2 and low level of TBXT at protein level (Fig 2D, S4B), a signature associated with NMPs (Attardi et al., 2018; Denham et al., 2015; Diaz-Cuadros et al., 2020; Edri et al., 2019; Faustino Martins et al., 2020; Gouti et al., 2014; Gouti et al., 2017; Lippmann et al., 2015; Metzis et al., 2018; Olivera-Martinez et al., 2012; Wymeersch et al., 2019). Overall, these results demonstrated that Wnt-induced hPSC-derived axial progenitors resemble axial progenitors of the mouse anterior caudal lateral epiblast that feed the elongation of the spinal cord (Edri et al., 2019; Gouti et al., 2014; Gouti et al., 2017; Henrique et al., 2015; Koch et al., 2017; Liu et al., 2001; Wymeersch et al., 2016).

**Figure 2.**
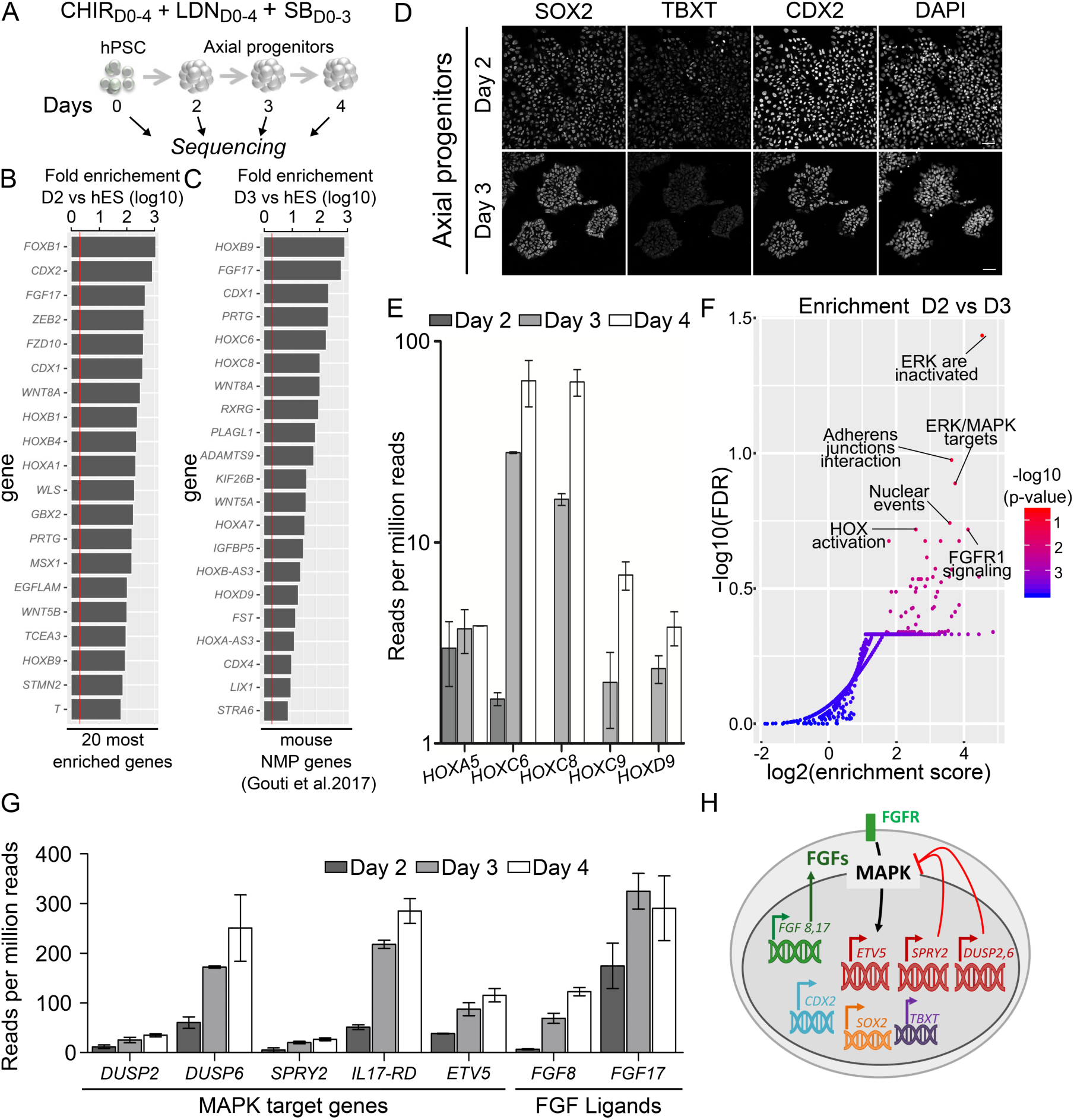
Temporal transcriptomic analysis of hPSC-derived axial progenitors. **(A)** Experimental design. **(B)** 20 most enriched genes in D2 progenitors versus hESC. Progenitors acquired a transcriptomic signature similar to the one of murine axial progenitors. **(C)** Fold enrichment in D3 progenitors versus hESC of the 20 most enriched neuromesodermal progenitor (NMP) genes defined in (Gouti et al., 2017). **(D)** Immunostaining for axial progenitor markers on hiPS-derived D2 and D3 progenitors. Cells co-express the three markers. Scale bars: 40 µm. **(E)** Temporal transcriptional changes in *HOX* genes coding for HOX regionally expressed in human MNs *in vivo*. **(F)** Reactome pathway analysis of the 232 genes upregulated 2 fold (p-value<0.05) between D3 and D2. Y axis: FDR = false discovery rate. X axis = enrichment score calculated for a given Reactome pathway. **(G)** Temporal evolution of upregulated MAPK target genes and FGF ligands in aging axial progenitors. ETV5 is a transcription factor activated by ERK1/2 MAPK in many other systems. The other genes encode feedback negative regulators of MAPK pathways. All data are shown as mean ± SD. **(H)** Schematic representation of the transcriptional and immunostaining analysis of day 2 and 3 progenitors.

Then, comparative analysis of the transcriptome of aging progenitors indicated a temporal collinear activation of *HOX* genes that display regionalized expression patterns in the spinal cord. At D2, the progenitors expressed *HOXA5* and low level of *HOXC6*, which belongs to the 3’ half of *HOX* complexes with anterior borders of expression in the anterior spinal cord (Fig. 1A, S1). On D3, *HOXC8* was well expressed and *HOXC9* expression had started, two genes belonging to the center part of *HOXC* complex with expression in the middle of the spinal cord (Fig. 1A, S1). Their expression further increased at day 4 (3.8 and 3.4-fold respectively) in progenitors that will give rise to HOXC8 and some HOXC9 MNs when exposed to RA/SAG (Fig. 1C-D). Non-expressed *HOX* genes corresponded to the 10 to 13 paralog groups, the latest and most caudally expressed *HOX* (Gaunt, 1991; Izpisúa-Belmonte et al., 1991; Philippidou and Dasen, 2013) and Fig. 1A, S1A,C). The temporality of the collinear activation in axial progenitors is thus in agreement with the generation of more caudal MNs when RA/SAG was delayed (Fig. 1). Hence, Wnt induced a temporal collinear activation of *HOX* genes, which parallels the change in rostro-caudal potential of the progenitors.

To define the pathways activated in parallel with the sequential induction of *HOX* genes, we performed a pathway analysis on the genes increased more than 2 fold (p<0.05) between D2 and D3. Among the 232 genes, we detected an enrichment for annotations associated with an activation of the Mitogen-activated protein kinases (MAPK) pathways due to a gradual increase in expression of typical MAPK target genes (*ETV5, DUSP4, DUSP6, IL17DR* or *SPRY2*) (Fig. 2F-H). MAPKs are classical mediators of FGF signaling (Lunn et al., 2007). In agreement, together with the rise in MAPK target genes we observed an increase in *FGF8* and *FGF17* expression, two secreted FGF ligands previously described to increase over time in the caudal epiblast of chick embryos (Fig. 2G-H) (Liu et al., 2001; Wymeersch et al., 2019).

Hence, the sequential collinear expression of *HOX* genes is paralleled by an increase in expression of *FGF* ligands and MAPK target genes within axial progenitors. As FGFs promote caudal *Hox* genes in other systems (Bel-Vialar et al., 2002; Dasen et al., 2003; Liu et al., 2001), we postulated that paracrine or autocrine FGF signaling might be triggering the sequential induction of *HOX* genes.

### FGF signaling is required for *HOX* sequential activation and caudal MN specification

We aimed at testing whether endogenous FGF signaling was necessary for the temporal shift in axial progenitor rostro-caudal potential and *HOX* gene induction. For that, we blocked FGF signaling in aging progenitors prior RA exposure and tested the impact on caudal *HOX* gene induction and MN subtype specification. We exposed progenitors from D3 to D7 to i) PD173074, a selective FGFR1/3 antagonist or ii) PD0325901 which inhibits the MAPK kinase MEK1/2 (Fig. 3A). The efficiency of MN generation was not impacted by the two inhibitors, even though we collected a reduced number of EBs in PD173074 condition (Fig. 3B). In control condition, RA/SAG at D7 induced MNs expressing HOXC8 (97.5%) and C9 (87.6%). In contrast, addition of the inhibitors at D3 while adding RA/SAG at D7 generated MNs with an anterior brachial identity (HOXC6). This more anterior identity was normally obtained when RA was added on early D3 progenitors showing that the inhibitors blocked the temporal change in rostro-caudal potential (Fig. 3A-C). We then tested whether the dissociation between the age of the progenitors and the rostro-caudal identity of progeny was linked to a stalled sequence of *HOX* gene induction in axial progenitors. We monitored the expression of *HOXC6, C8* and *C9* mRNAs following MEK1/2 inhibition at D3. While in control condition, the three genes increased over time, MEK inhibition blocked this increase at D4 or D5 (Fig. 3D). Hence, inhibition of FGF signaling in aging axial progenitors blocks *HOX* temporal induction and caudal cell type specification. Our transcriptomic analysis indicated a temporal increase in *FGF8* and *FGF17* expression suggesting that an increase in FGF concentration and/or duration of exposure might be pacing the *HOX* clock.

**Figure 3.**
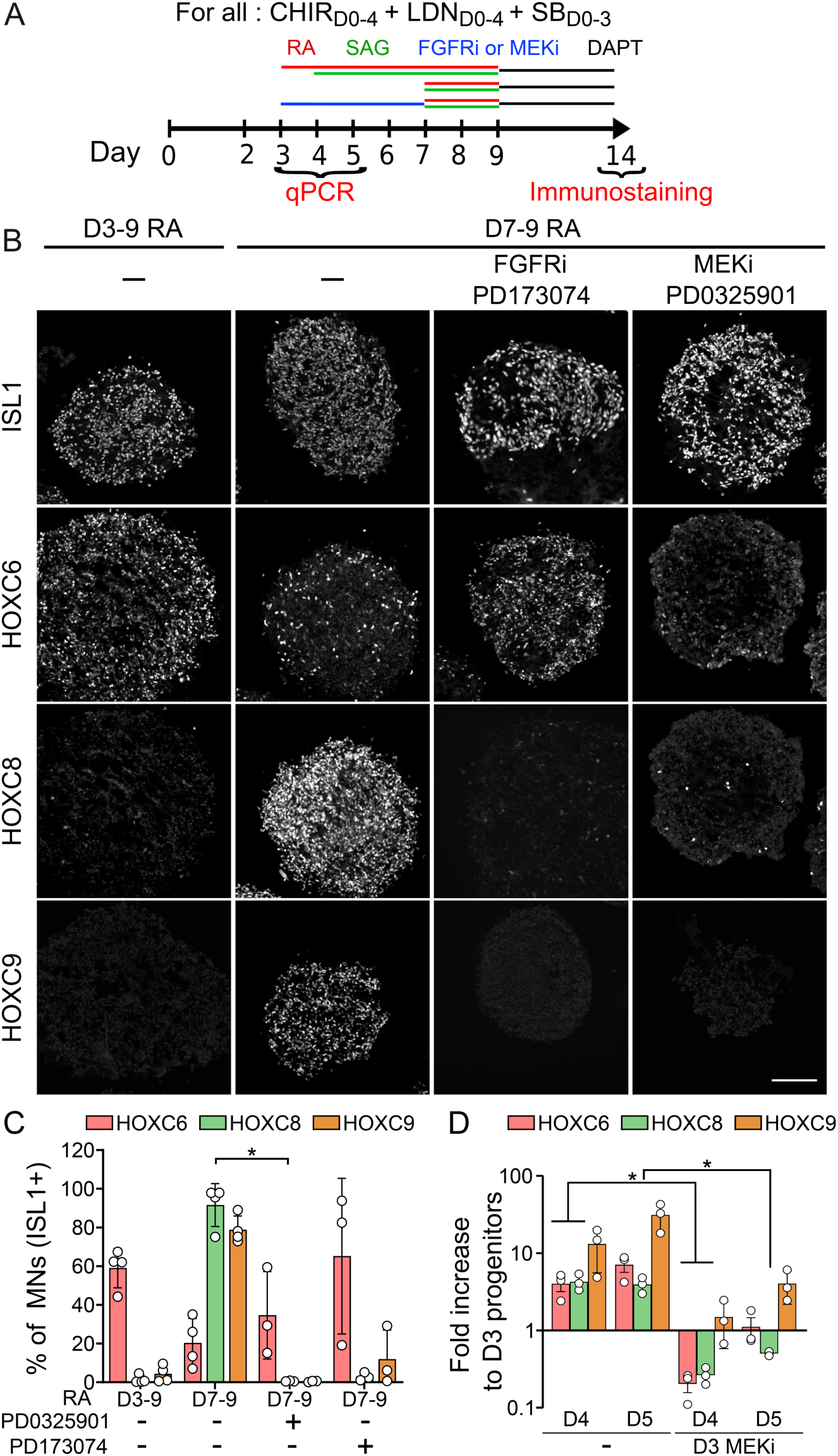
FGF pathway inhibition stalls *HOX* temporal induction and caudal MN specification. **(A)** Differentiation conditions. PD173074 (FGFR1-3 inhibitor) or PD032590 (MEK1/2 inhibitor) were added at D3 up to D7. **(B)** Proportion of MNs (ISL1+ cells) in the different conditions (mean ± SD). **(C)** Immunostaining for ISL1 (MNs) and HOX transcription factors on cryostat sections of embryoid bodies on day 14 of differentiation. MEK and FGFR inhibitors prevent the specification of HOXC8 and HOXC9 MNs. Instead, HOXA5, HOXC6 MNs are generated. Scale bar: 100µm. **(C)** Quantification of HOXC6 and HOXC9 MNs on day 14 of differentiation. **(D)** Real time PCR analysis of *HOX* mRNAs in progenitors at D3, D4 and D5 of differentiation. MEK and FGFR inhibitors prevent the temporal increase in caudal *HOX* expression. Data are shown as mean ± SD; Each circle is an independent experiment. * if P ≤ 0.05, ** if P ≤ 0.01 and *** if P ≤ 0.001.

### FGF level paces the *HOX* clock and MN subtype specification

To test whether the level of environmental FGF paces the *HOX* clock, we exposed early D3 progenitors, to RA/SAG together with FGF2 at different concentrations and for different durations. In all conditions that received FGF2, more caudal MN identities were induced without the need of delaying RA addition (Fig. 4A-D, S3). Then, the extent of caudalization varied with FGF parameters. Exposing progenitors from D3 to D9 to increasing concentrations of FGF2 induced progressively more caudal MN subtypes. Caudal brachial HOXC8 MNs were induced at 15 ng/ml (68.2%, 8.6-fold increase) and HOXC9+/HOXC6-thoracic MNs at 60 ng/ml (58.9 %, 82.5-fold increase) while 120 ng/ml reduced slightly more HOXC6 MNs (Fig. 4A-D, S6A-H). Similar results were obtained with FGF8 (Fig. S3B). FGFs acted directly and rapidly on axial progenitors as addition of 120 ng/ml FGF2 for 24h or 48h induced MNs of a caudal brachial or mid-thoracic identity (49.7% HOXC6, 79.4% HOXC8, 14.7% HOXC9, 47.9% FOXP1^high^/SCIP for 24h) (7.5% HOXC6, 76.4% HOXC9, 13.2 % FOXP1 ^high^/SCIP for 48h) (Fig. 4A-C, S3C-H). To determine whether this caudalizing effect was underlined by an accelerated induction of brachial and thoracic *HOX* genes we performed real-time PCR analysis 24h and 48h post FGF2 treatment. We observed a precocious increase in *HOXC8, HOXC9, HOXD9* and *HOXC10* expression compared to RA/SAG controls (Fig. 4F).

**Figure 4.**
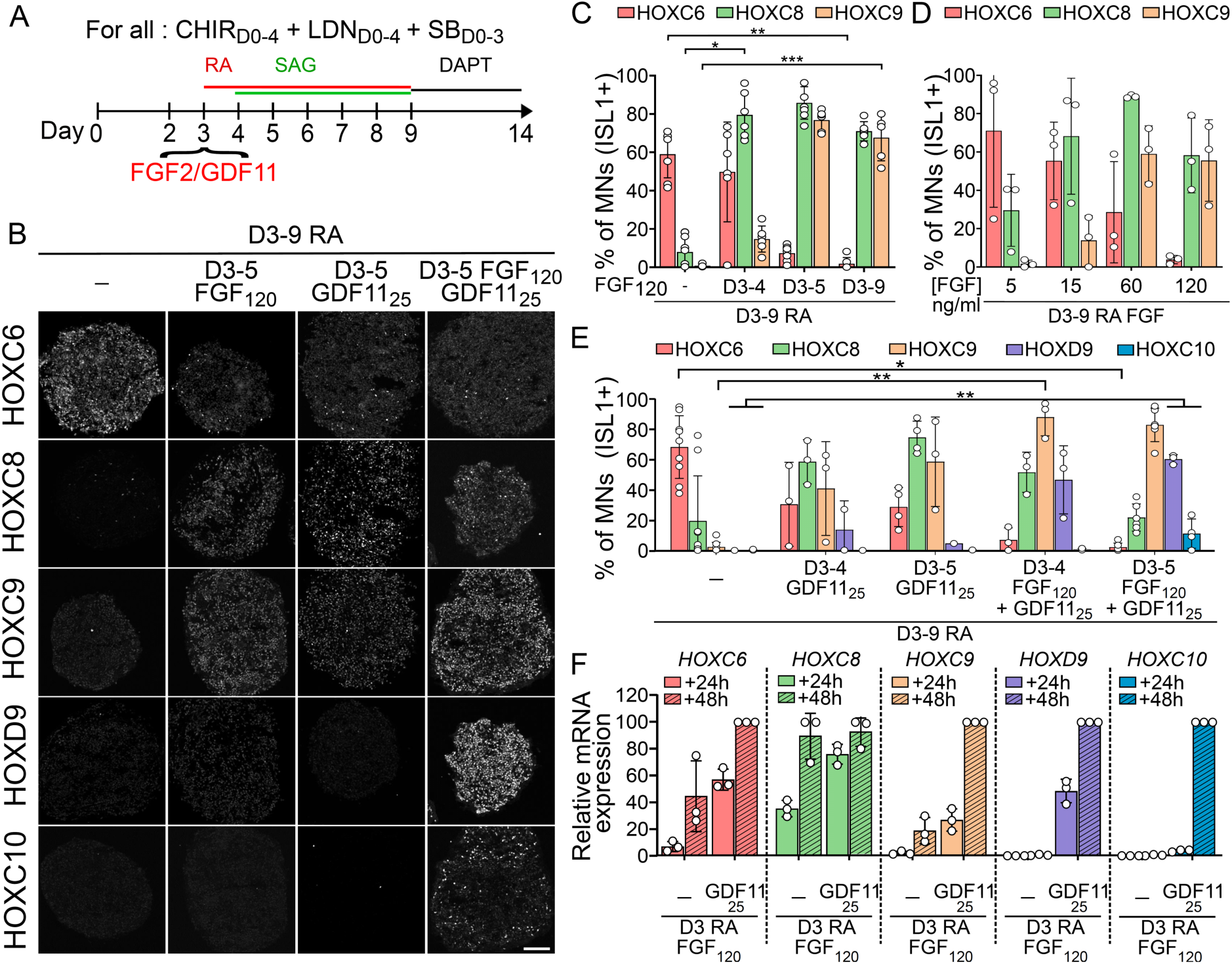
Dynamic pacing of *HOX* induction and specification of discrete MN subtypes upon changes in FGF2 and GDF11 levels. **(A)** Differentiation conditions. Extrinsic cues, FGF2, GDF11, or FGF2+GDF11 were added on day 3 of differentiation at various concentrations or for different durations **(B)** Immunostaining for HOX on cryostat sections of embryoid bodies on day 14 of differentiation. FGF2, GDF11 and FGF2/GDF11 induce more caudal MN subtypes. Scale bar: 100µm. **(C, D, E)** Proportion of MNs (ISL1+ cells) expressing the indicated markers. The effect of the duration of FGF2 treatment (C), FGF2 concentration (D) and duration of GDF11 or FGF2+GDF11 (E) were monitored. **(F)** Real time quantitative PCR analysis of the expression of *HOX* genes regionally expressed in human MNs *in vivo. HOX* mRNAs were monitored at D4 (24h post-treatment) and D5 (48h post-treatment) upon addition of FGF2, GDF11 or a combination of FGF2 (120 ng/ml) together with GDF11 (25 ng/ml). Data is normalized at each time point to the control condition treated with only RA at day 3. Data are shown as mean ± SD. Each circle is an independent experiment. * if P ≤ 0.05, ** if P ≤ 0.01 and *** if P ≤ 0.001. See also Fig. S3

Hence, a precocious increase in FGF signaling accelerates the induction of caudal *HOX* genes in early axial progenitors resulting in the specification of more caudal cell types within the same differentiation timeline (14 days). These results demonstrate that the levels or duration of FGF signaling dynamically pace the tempo of *HOX* collinear activation during human axial progenitor differentiation.

### FGF and GDF11 synergize to further accelerate the *HOX* clock

FGF2 accelerated *HOX* induction to generate MNs up to the mid-thoracic level suggesting that early axial progenitors might not be competent to generate more caudal segments. Alternatively, other extrinsic factors might be required to further accelerate the induction of *HOX* genes and promote more caudal identities. GDF11 is a member of the TGFβ family implicated in the control of axial elongation, MN subtype specification and is required for the expression of the most 5’ *HOX* gene starting from the 10 paralogs *in viv*o and *in vitro* in late NMP-like cells (Aires et al., 2019; Gaunt et al., 2013; Lippmann et al., 2015; Liu, 2006; Liu et al., 2001; McPherron et al., 1999; Peljto et al., 2010). We exposed D3 early progenitors to a combination of RA and GDF11 (25 ng/ml) for 24h or 48h. 24h GDF11 induced caudal brachial and anterior thoracic MNs (58.7 % HOXC8, 41.0 % HOXC9); the latest category increasing after 48h (Fig. 4A, E, S4A). However, as with FGF2, none of the more caudal identities were observed. In chick, exposure of spinal cord explants to combination of FGF2 and GDF11 promoted more caudal MNs than the two factors separately (Liu et al., 2001). We thus tested whether this combination might accelerate the clock and promote caudal thoracic or lumbar identities. 24h after FGF2/GDF11 treatment, *HOXC9, HOXD9* and *HOXC10 mRNAs* were strongly induced (respectively 65.4, 2774.77 and 3329.44-fold compared to controls) and further increased after 48h (Fig. 4F). In agreement, a 24h (D3-D4) treatment induced, on day 14, caudal thoracic MNs (88.2 % HOXC9 and 46.7% HOXD9) and lumbar HOXC10 MNs after 48h of exposure (11.1%) (Fig. 4B, E, S4B-E).

Hence, axial progenitors are competent to produce the different spinal rostro-caudal identities from brachial to lumbar. Changes in parameters of exposure to FGFs and GDF11 (concentrations, durations and combinations) dynamically pace the speed at which the *HOX* clock proceed resulting in the specification of early or later born MN subtypes within the same timeline of differentiation.

## DISCUSSION

Overall, we report the selective generation of axial progenitors from hPSCs demonstrated by combination of specific markers, detailed transcriptomic analysis and their ability to generate cell types found at distinct rostro-caudal levels. Importantly, we observed line to line differences in the optimal concentration of CHIR necessary to potently induce an axial progenitor state for the subsequent reliable induction of caudal cell types. With efficient access to axial progenitors we then show that changes in the concentration, duration and combination of the caudalizing factors FGFs and GDF11 control the speed at which the temporal collinear activation of *HOX* genes occurs. The pace of the *HOX* clock is thus dynamically encoded by the parameters of exposure to extrinsic cues. The sequential changes in chromatin structure occurring along *HOX* complexes during their activation do not appear to act as an intrinsic timer actuated and stopped by extrinsic factors (Bel-Vialar et al., 2002; Del Corral and Storey, 2004; Kimelman and Martin, 2012; Lippmann et al., 2015; Mazzoni et al., 2013; Narendra et al., 2015; Noordermeer et al., 2011; Noordermeer et al., 2014; Soshnikova and Duboule, 2009; Tschopp et al., 2009). Our results might provide a molecular basis for the observation that heterochronic grafting of “old” axial progenitors to a “younger” caudal stem zone reverted their *HOX* profile to the “young” one (McGrew et al., 2008; Wymeersch et al., 2019). The pacing of the *HOX* clock by secreted factors might ensure a community effect to synchronize *HOX* expression between neighboring progenitors which could allow the emergence of expression domains at tissue level (Durston, 2019). Furthermore, FGFs and GDFs also control the maintenance of the stem cell pool and axial elongation (Aires et al., 2019; Boulet and Capecchi, 2012; Jurberg et al., 2013; Mallo et al., 2009; McPherron et al., 1999). A common mechanism to control rostro-caudal extension of the body axis together with *Hox* gene induction would be a parsimonious way to couple morphogenesis and patterning (Denans et al., 2015; Young et al., 2009). In a bioengineering perspective, *HOX* clock extrinsic control provides a simple mean to engineer cell types of defined “rostro-caudal” identities from PSCs. It shortens temporal requirements for the generation of cells born at different time of development during axial elongation. This allows the synchronous engineering with an unprecedented efficiency and precision of human MN subtypes with defined rostro-caudal identities. In particular, we provide the first evidence for the generation of putative hand-controlling MNs (Mendelsohn et al., 2017). MN subtypes display differential vulnerabilities in disease and in spinal injuries. Our results will provide access to a long awaited resource for the modeling of these incurable diseases and more controlled source of cells for putative cell therapy approaches. (Abati et al., 2019; An et al., 2019; Baloh et al., 2018; Nijssen et al., 2017; Ragagnin et al., 2019; Sances et al., 2016; Steinbeck and Studer, 2015; Tung et al., 2019). As HOX play a central role in instructing cell diversification in the three lineages, controlled manipulation of the *HOX* clock has a great potential for cell engineering besides MN subtypes. Our strategy is likely expandable to other axial stem cell derivatives such as paraxial mesoderm that generate the different muscles of the body with clear applications for the modeling of neuromuscular diseases (Bakooshli et al., 2019; Diaz-Cuadros et al., 2020; Faustino Martins et al., 2020; Frith et al., 2018; Machado et al., 2019; Matsuda et al., 2020; Osaki et al., 2020; Pourquié et al., 2018; Steinbeck et al., 2015; Verrier et al., 2018). More broadly, the temporal generation of distinct types of neurons or glia from the same progenitor domain is a widely used strategy to increase cell diversity in the nervous system (Dias et al., 2014; Kohwi and Doe, 2013; Oberst et al., 2019a; Rossi et al., 2017). Extrinsic cues play important roles in the unfolding of these temporal sequences (Kawaguchi, 2019; Oberst et al., 2019a; Oberst et al., 2019b; Syed et al., 2017; Tiberi et al., 2012). Extrinsic manipulation of the temporality of these lineages should improve the generation of early and late-born cells for both basic research, disease modeling and cell therapy.

## MATERIALS and METHODS

### Human embryonic spinal cord histology

Human fetal embryo of 6.3 weeks of gestation were obtained from pregnant women referred to the Department of Gynecology and Obstetrics at the Antoine Béclère hospital (Clamart, France) for legally induced abortions in the first trimester of pregnancy as previously described (Lambrot et al., 2006). All women provided written informed consent for scientific use of the fetal tissues. None of the abortions were due to fetal abnormality. The fetal age was determined by measuring the length of limbs and feet (Evtouchenko et al., 1996). The project was approved by the local Medical Ethics Committee and by the French Biomedicine Agency (reference number PFS 12– 002). Alternatively, Human embryonic spinal cords (n=2) at stage 7.5 weeks of gestation were collected in accordance with the national guidelines of the United States (National Institutes of Health, U.S. Food and Drug Administration) and the State of New York and under Columbia University institutionally approved ethical guidelines relating to anonymous tissue. The material was obtained after elective abortions and was classified on the basis of external morphology according to the Carnegie stages. Gestational age was determined by last menstrual period of the patient or by ultrasound, if the ultrasound estimate differed by 1 week as indicated by the obstetrician. In all cases, the spinal cord was removed as intact as possible before fixation with fresh, cold 4% PFA for 1.5 h on ice washed abundantly with PBS and then cryoprotected overnight in 30% sucrose. Post-fixation, the cord was measured and cut into anatomical sections to accommodate embedding in OCT Compound (Leica) and stored at −80°C before cutting on a cryostat. Sections (16 µm) were cut along the full length of the cord.

### Human pluripotent stem cell lines

Human SA001 embryonic stem cell (ESC) line (male, RRID: CVCL_B347) was obtained from Cellectis and used accordingly to the French current legislation (Agency of Biomedicine, authorization number AFSB1530532S). Induced pluripotent stem cell (iPSC) line WTSIi002 (male, RRID: CVCL_AH30, alternative name HPSI0913i-eika_2) and WTSIi008-A (male, RRID: CVCL_AH70, alternative name HPSI1013i-kuxp_1) were obtained from the European Bank for Pluripotent Stem Cells (EBISC). Experiments with iPSCs were approved by relevant ethic committees (declaration DC-2015-2559). All PSC lines were cultured at 37 °C on Matrigel (Corning) in mTSER1 medium (Stem Cell Technologies) and amplified using EDTA (Life Technologies) clump-based passaging. They were tested for potential mycoplasma contamination every other week (MycoAlertTM Mycoplasma Detection Kit, Lonza, LT07-118). No contamination was detected during the study. PSC were thawed in presence of Y-27632 (10µM, Stemgent or Stem Cell Technologies) and the culture media was changed every day.

### Human pluripotent stem cell differentiation

Human PSC embryoid body-based differentiation was performed as previously described (Maury et al., 2015). hPSC were dissociated with cold Accutase (Life Technologies) for 3 to 5 min at 37°C and resuspended in differentiation medium N2B27 (Advanced DMEM F12, Neurobasal vol:vol (Life Technologies)), supplemented with N2 (Life Technologies), B27 without Vitamin A (Life technologies), penicillin/streptomycin 1%, β-Mercaptoéthanol 0.1% (Life Technologies), with Y-27632 (10µM, Stemgent or Stem Cell Technologies), CHIR-99021 (3 µM or 4 µM Selleckchem) SB431542 (20 µM, Selleckchem) and LDN 193189 (0.1 µM, Selleckchem). 2×10^5^ cells ml^-1^ were seeded in ultra-low attachment 6 well plates (Corning) to form embryoid bodies (EBs). All conditions of differentiation received the same medium at day 2 and at day 3 but SB431542 was removed at day 3. Then, the differentiation proceeded according to the schemas presented above the figures. SAG (Smoothened Agonist, Merck millipore), FGF2 (Recombinant Human FGF basic, Peprotech), RA (Sigma-Aldrich), GDF11 (Recombinant Human/Murine/Rat GDF11, Peprotech), PD0325901 (Selleckchem), PD173074, SCH772984 (Selleckchem), DAPT (Stemgent) were added at indicated time points. For concentrations see Table S2. Media were changed every other day unless specified.

### Embryoid body processing for immunostaining

EBs were collected, rinsed with PBS then fixed with 4% PFA for 5 min at 4°C and rinsed with PBS 3 times for 5 min. EBs were cryoprotected with 30% sucrose and embedded in OCT (Leica) prior sectioning with a cryostat. Alternatively, day 2, 3 and 4 progenitors were plated on Matrigel (Corning, diluted according to manufacturer recommendation) coated coverslips and let to adhere between for 30 min prior fixation with 4% PFA for 5 min at 4°C.

### Immunostaining

All immunostainings were performed as follow: cells or sections were incubated with a saturation solution (PBS/FBS 10 %/0.2% Triton) for 10 minutes. Primary antibodies (Table S3) were diluted in staining solution (PBS/2% FBS/0.2% Triton) and incubated overnight at 4°C in a humidified chamber. Following 4 PBS washes (10 min each), secondary antibodies (Alexa488, Alexa568 and Alexa647, Life Technologies, 1: 1,000) were added for 1h at RT. After 3 PBS washes, DAPI was added on the cells (Invitrogen, 1: 3,000) for 5 min. Cells or slices were then mounted in Fluoromount (Sigma Aldrich or Cliniscience).

### Image acquisition

Samples were visualized and imaged using either a ZEISS LSM 880 Confocal Laser Scanning Microscope (Carl Zeiss Microscopy) controlled by Zen black software (Zeiss), a confocal microscope TCS SP5 II (Leica) or a DM6000 microscope (Leica) equiped with CoolSNAP EZ CDD camera, controlled by MetaMorph software (MetaMorph Inc). Alternatively, images were acquired using the automated microscope Cell Discoverer 7 (Zeiss), equipped with an Axiocam 506m camera, with Zen black software (Zeiss).

### Quantitative RT-PCR analysis

Total RNA were extracted (RNAeasy Plus Mini Kit, Qiagen) and cDNA synthesized using SuperScript III (Invitrogen). Quantitative real-time PCR was performed using a 7900HT fast real time PCR system (Applied Biosystems) with Sybr Green PCR Master Mix (Applied Biosystems). Alternatively performed using QuantStudio 5 Real-Time PCR System (Thermofisher Scientific) and a mix with qPCR Brilliant II SYBR MM with low ROX (Agilent). Primers are listed in Supplementary Table 3. All expression data were normalized to *Cyclophilin* A mRNA. All analyses were performed with three technical replicates per plate. Relative expression levels were determined by calculating 2−ΔΔCt.

### Transcriptomic analysis

hESC were collected post dissociation and prior exposure to differentiation medium. Progenitors were collected on day 2, day 3 and day 4 of differentiation. For each of the 8 samples, total RNA were extracted then reverse transcribed using the Ion AmpliSeq Transcriptome Human Gene Expression kit (Thermofisher Scientific). The cDNA libraries were amplified and barcoded using Ion AmpliSeq Transcriptome Human Gene Expression core panel and Ion Xpress Barcode Adapter (Thermofisher Scientific). The amplicons were quantified using Agilent High Sensitivity DNA kit before the samples were pooled in sets of eight. Emulsion PCR and Enrichment was performed on the Ion OT2 system Instrument using the Ion PI Hi-Q OT2 200 kit (Thermofisher Scientific). Samples were loaded on an Ion PI v3 Chip and sequenced on the Ion Proton System using Ion PI Hi-Q sequencing 200 kit chemistry (200 bp read length; Thermofisher Scientific). The Ion Proton reads (FASTQ files) were imported into the RNA-seq pipeline of Partek Flow software (v6 Partek Inc) using hg19 as a reference genome. The number of reads per sample was ranging from 7.5 million to 12 million reads. To determine genes that are differentially expressed between groups mapped reads were quantified using Partek E/M algorithm normalized by the Total count/sample (the resulting counts represent the gene expression levels on reads/millions for over 20,800 different genes present in the AmpliSeq Human Gene Expression panel). The evaluation of the differential expression between two conditions was performed using the EdgeR package under R. Pathway enrichment analyses were performed on upregulated genes (FC⩾2.0, p-value<0.05) between two time points by interrogating Reactome database. Significant enrichments were calculated using hypergeometrical test and Benjamini-Hochberg correction for multiple comparisons. The enrichment score was calculated as described in (Wang et al., 2017). The normalized transcriptomic data are provided in Table S1. Raw data are available upon request.

### Quantification and statistical analysis

All statistics were computed using Graphpad Prism software. One-way analysis of variance (ANOVA) with a Kruskall-Wallis post hoc analysis were performed following normality tests provided by Prism. Number of n, dispersion measures and P-values are indicated in figure legends. In all figures, n are independent differentiations started from independent newly thawed hPSCs vials. For each condition at least 4 independent EBs were imaged in which all the cells were quantified by automated image analysis. The images were exported and saved as TIFF with Fiji if needed (Schindelin et al., 2012). Quantitative analyses on images were performed using the CellProfiler software (Carpenter et al., 2006) (Broad Institute open source at www.cellprofiler.org). DAPI stained nucleus were segmented into primary objects using CellProfiler segmentation pipeline and the nuclear mask was used to define objects on the target channels. The threshold to define positive nuclei for a given target was obtained using EBs’ section negative for the target of interest. All images across conditions were then automatically analyzed in batch ensuring unbiased analysis. The analyses of the FOXP1 and SCIP immunostaining intensity were performed combining CellProfiler with the software FCS express 7 (DeNovo Software, Glendale, CA, USA). Nuclei were segmented into primary objects as described above and FOXP1 and SCIP fluorescence intensities were calculated in each primary object. Fluorescence intensity plots for FOXP1 and SCIP were then generated using FCS express 7 software to visualize the intensity levels of the different markers for each individual cells and determine the percentage of cells above a given threshold. Cell profiler pipelines for quantification are available upon request.

## ACKNOWLEDGEMENTS AND FUNDING

This project was supported by grants from AVENIR/ATIP, Association Française contre les Myopathies (AFM) and Laboratoire d’Excellence (Labex) Biopsy and Revive (Investissement d’Avenir (ANR-10-LABX-73 and 11-LABX-0035, respectively). V.M has received fellowships from Domaine d’Investissement Majeur (DIM) “Biothérapies” of the region Ile-de-France and from the Fondation pour la Recherche Médicale (FRM). C.V. is supported by a PhD fellowship of the French Ministry of Research. V.R. and S.N salaries are funded by the INSERM. We thank Frédéric Causeret for providing sections of mouse embryos. We thank Susan Morton, Thomas Jessell and Jeremy Dasen for the generous gift of antibodies. We thank Fréderic Causeret, Pascale Gilardi, Francois Giudicelli and Esteban Mazzoni for critical reading of the manuscript. Imaging experiments were carried out at the Imaging platform of the Institut du Fer à Moulin and at I-STEM.

## AUTHOR CONTRIBUTIONS

S.N conceived the project. V.M, C.V, S.N, C.M, designed experiments. V.M, C.V, S.G, performed most experiments and received assistance with pluripotent stem cell culture from L.L and A.T and with data processing and analyses from V.R, R.R, M.D, M.J. RNA sequencing: M.J. Human spinal cord analysis: V.M, N.N, V.R.F, G.C, MWM, HW. Project Administration: S.N, C.M. Supervision: S.N, C.M, H.W, V.RF. Writing: - Original Draft: S.N V.M-Review & Editing: all authors

## MATERIALS AVAILABILITY

The normalized transcriptomic data are provided in Table S1. FASTQ files will available in GEO. Cell profiler pipelines for quantification are available upon request. Further information and requests for resources and reagents should be directed to

## SUPPLEMENTARY FIGURES AND TABLES

**Figure S1.**
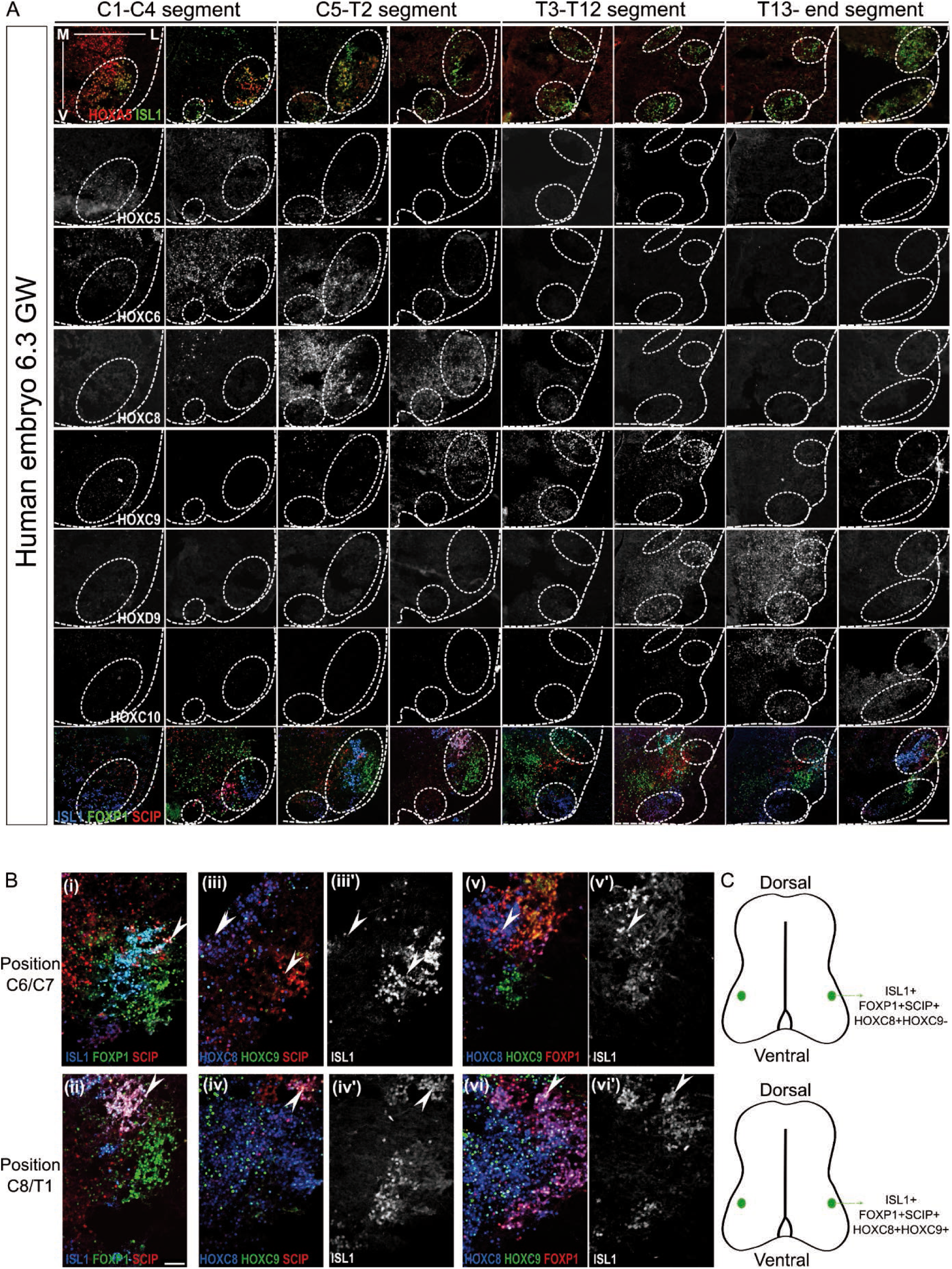
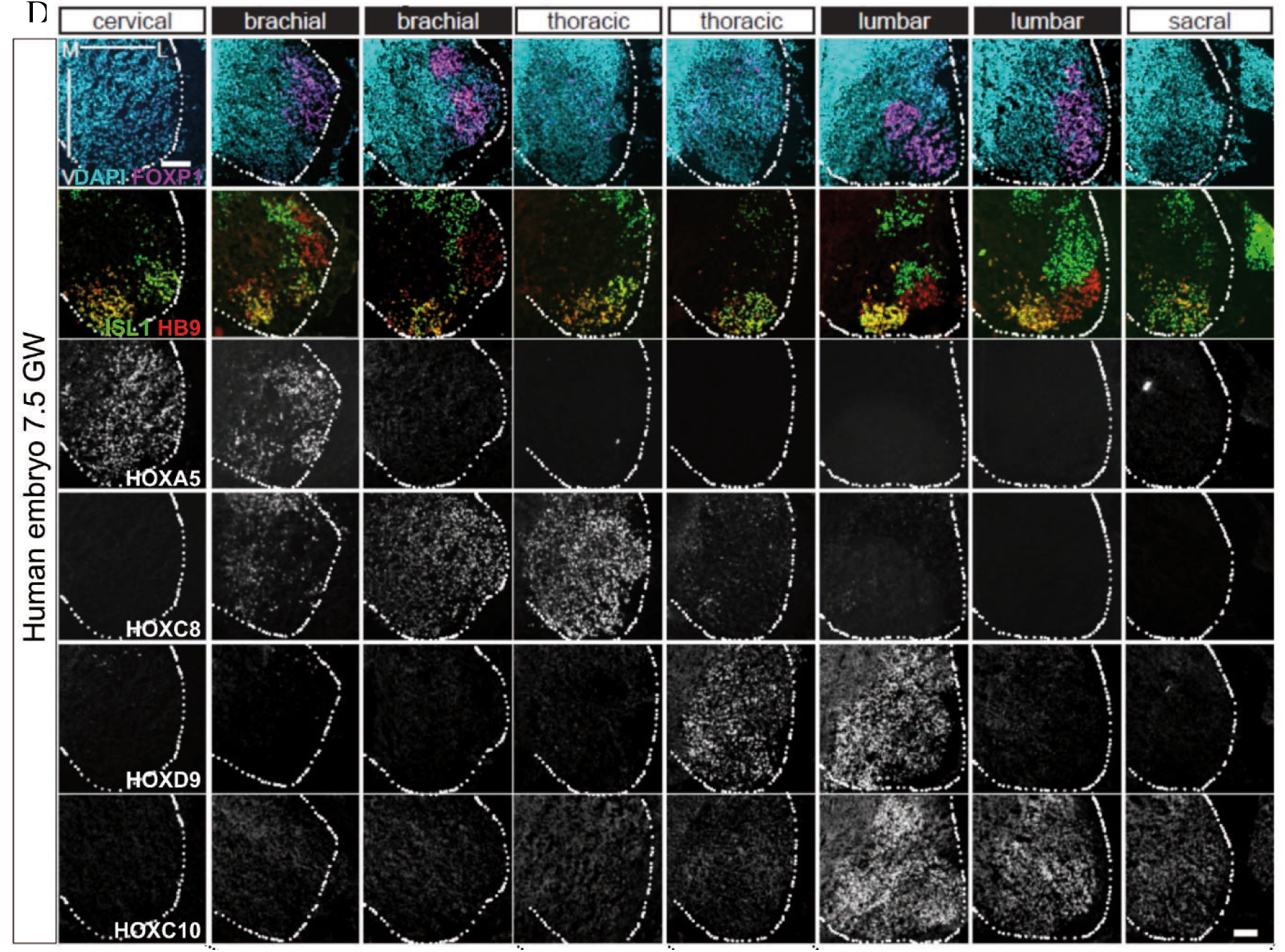
Rostro-caudal distribution of motor neuron subtypes in human embryonic spinal cords. *Data used to construct Figure 1A*. Immunostaining on transverse sections of human embryonic spinal cords at gestation weeks (GW) 6.3 **(A, B)** and 7.5 **(D)** to define HOX expression profiles in spinal motor neurons and localization of MN subtypes. MNs organized in motor columns that innervate distinct muscle groups were identified by the expression of ISL1 or HB9. External dotted lines in A and D mark the side of the ventral horn. M: medial L: lateral V: Ventral. **(C)** Schematic representation of the FOXP1/SCIP/ISL1/HOXC8 and FOXP1/SCIP/ISL1/HOXC9 MNs in brachial and thoracic regions. Scale bars: 100μm HOXA/C5 MNs are localized in cervical regions, while brachial MNs expressed HOXC6, thoracic HOXC9 and lumbar HOXC10. HOXC8 is expressed in caudal brachial MNs with HOXC6 and in the anterior thoracic domain with HOXC9. HOXD9 identified the caudal thoracic and the anterior lumbar MNs. **(A, B**,**D)** As in mouse and chick, FOXP1^high^ MNs at brachial and anterior thoracic levels form a lateral motor column (LMC) in a region where limb-innervating MNs are located (Dasen et al., 2008; Routal and Pal, 1999). In the thoracic mouse spinal cord, FOXP1^low^ MNs form the most dorsal motor column containing preganglionic sympathetic MNs (Dasen et al., 2008) also observed here. **In A and B**, FOXP1/SCIP/ISL1 MNs are observed in the caudal brachial and anterior thoracic spinal cord which in mouse identify MNs innervating respectively forearm and forepaw muscles (Catela et al., 2016; Dasen et al., 2003; Jung et al., 2010). In mouse, brachial SCIP MNs that express Hoxc8 innervate forearm muscles while anterior thoracic SCIP MNs co-express Hoxc8 and Hoxc9 and innervate the digit controlling muscles (Mendelsohn et al., 2017). These MN subpopulations are conserved in human: **in B iii) and v)** arrows point to the presence of HOXC9 negative FOXP1/HOXC8/ISL1 positive MNs while in **B iv) and vi)** SCIP/ISL1 or FOXP1/ISL1MNs in anterior thoracic domains coexpress HOXC8 and HOXC9 which might thus correspond to hand-innervating MNs. In mouse, SCIP^high^ MNs at cervical level are phrenic innervating MNs (Machado et al., 2014; Philippidou et al., 2012). SCIP^low^ MNs along the spinal cord are a subset of axial innervating muscles (Rousso et al., 2008). These populations are conserved in human.

**Figure S2.**
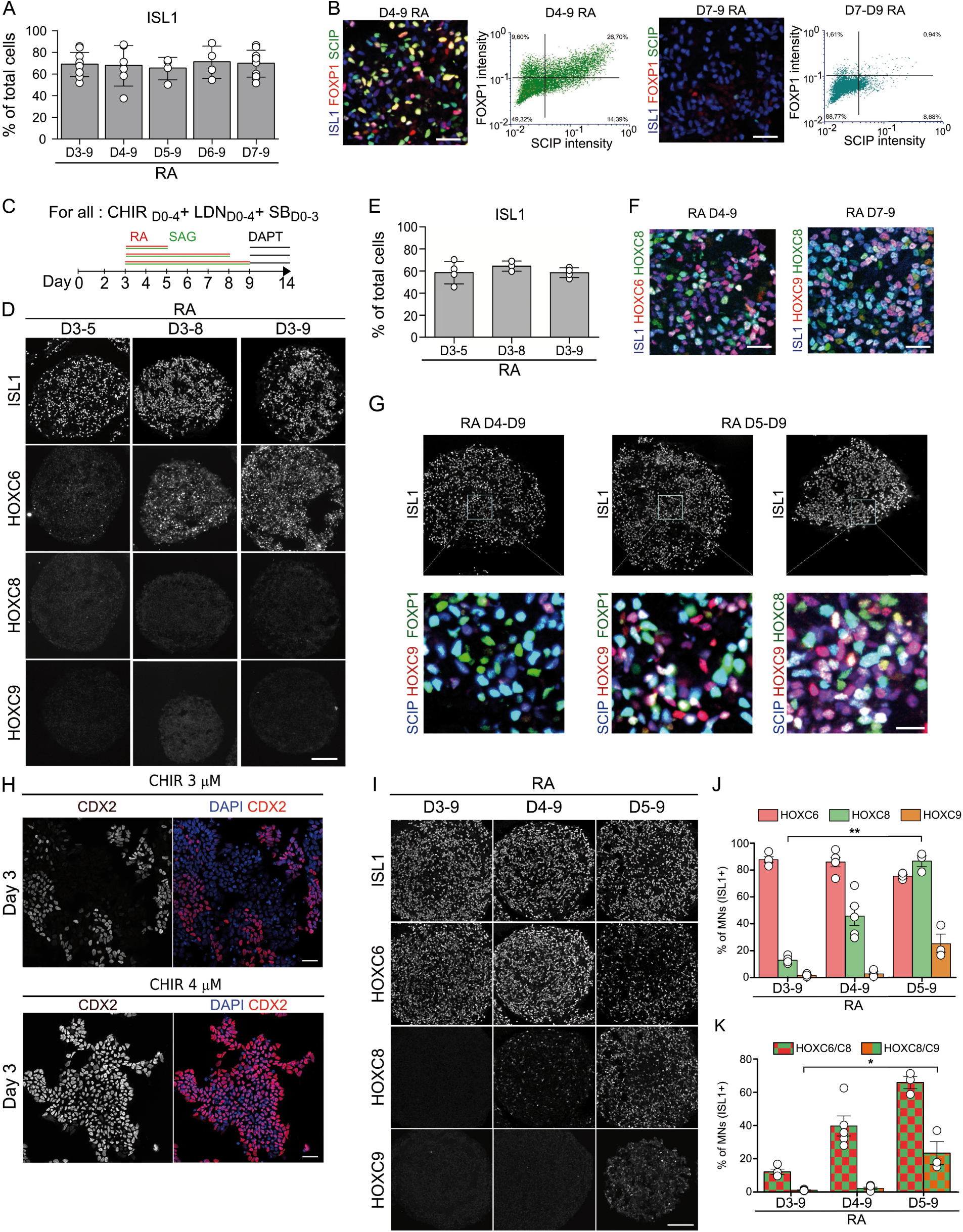
The timing of exposure to RA and not the exposure duration controls the specification of caudal motor neurons from hESC and hiPSC. **(A)-** Proportion of MNs (ISL1+ cells, mean ± SD) generated on day 14 of differentiation when modulating the day at which RA/SAG is added. **(B)** Quantification of FOXP1^high^ and FOXP1^high^/SCIP MNs. Intensity of FOXP1 and SCIP staining in all ISL1+ cells were extracted using Cell profiler. Intensity plots were generated for all conditions. FOXP1 and SCIP weak or negative cells were defined on the D7-D9 RA condition. **(C-E)** Characterization on day 14 of differentiation of the effect of varying RA/SAG duration of exposure on MN subtypes identity. (C) Differentiation conditions. (D) Immunostaining on cryostat sections of EBs for ISL1 and HOXs showing the absence of caudal HOXs when RA is added on day 3 and the duration of exposure modulated. (E) Proportion of MNs (ISL1+ cells, mean ± SD) generated on day 14 in the three tested conditions. **(F**) Immunostaining on cryostat sections of D14 EBs for the indicated markers quantified in Fig. 2F. **(G)** Immunostaining on cryostat sections of EBs showing the presence of HOXC8/HOXC9/SCIP/ISL1 MNs and HOXC9/SCIP/FOXP1/ISL1 MNs that correspond to forepaw innervating-MNs in mouse and might represent hand innervating MNs in human. Scale bar: 15µm. **(H)** Immunostaining on D3 EBs plated on matrigel coated coverslips. EBs were exposed either to CHIR 3 or 4 µM on day 0 of differentiation together with LDN and SB. Scale bar: 40 μm. **(I)** Immunostaining on cryostat sections of D14 EBs for HOXs showing the generation of HOXC8 and then HOXC9 MNs when addition of RA is delayed. Scale bar: 100 μm. **(J)** Quantification of indicated markers (mean ± SD, each circle = 1 independent experiment).

**Figure S3.**
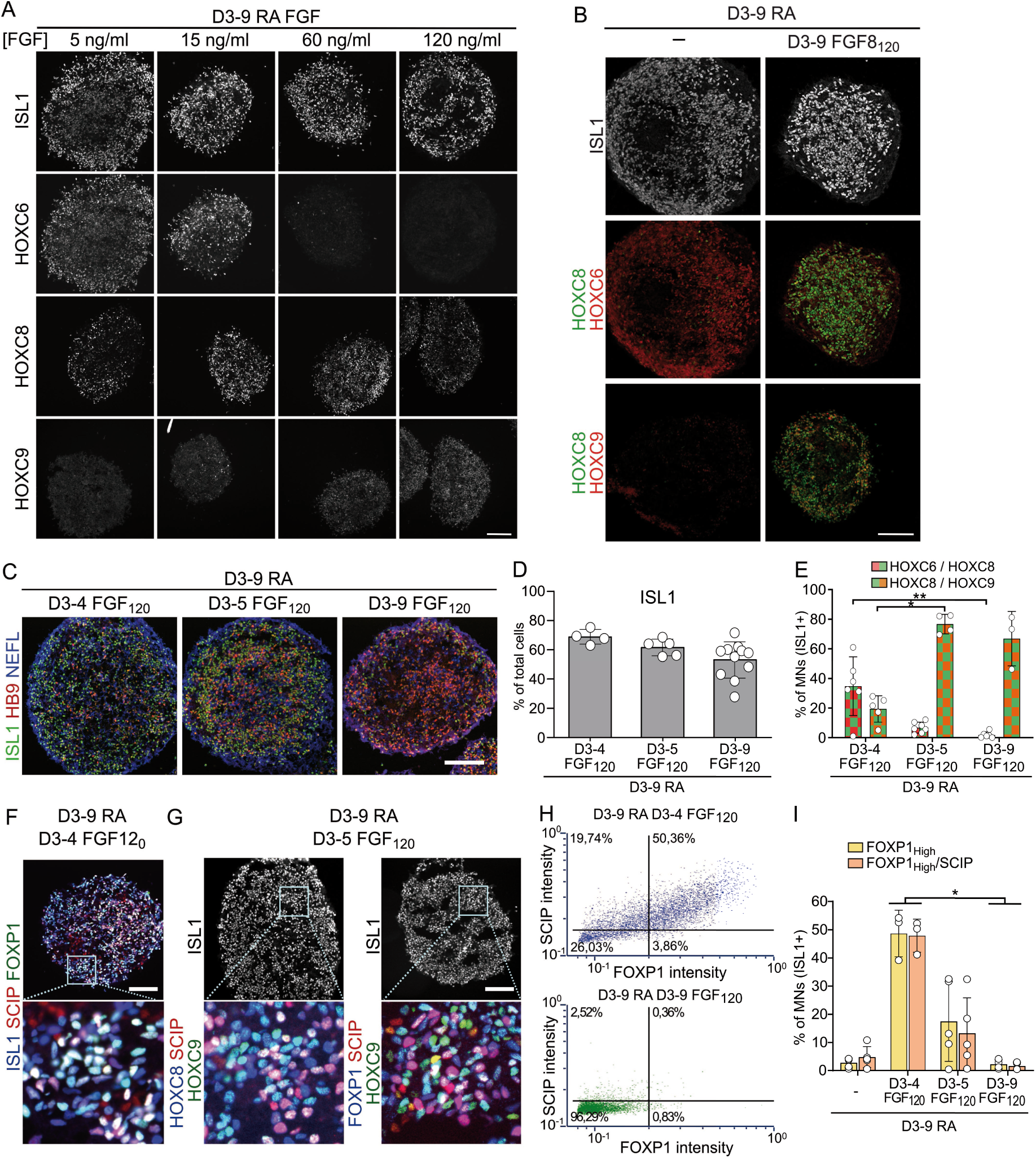
Concentration and duration of FGF control motor neuron subtype identity. **(A)** Immunostaining on EB cryostat sections at D14. Staining for ISL1, HOXC6, HOXC8 and HOXC9 to test the effect of changes in FGF2 concentration on MN subtype identity. **(B)** Immunostaining for ISL1, HOXC6, HOXC8, HOXC9 in control condition and with 120 ng/ml FGF8 (D3-9). **(C-I)** Characterization of MN generation and MN subtype identity at D14 following changes in FGF2 duration of exposure - FGF2 was added at D3 for different durations. (C) Immunostaining on cryostat sections of EBs for ISL1, HB9 and NEFL to assess MN specification. (D) Proportion of MNs at D14. (E) Quantification of HOXC6/HOX8/ISL1 and HOXC9/HOXC8/ISL1 co-expression. (F-G) Immunostaining for ISL1, SCIP and FOXP1 (E) or HOXC8/HOXC9/SCIP/ISL1 (F) aiming at characterizing the specification of MNs acquiring the identity of limb or digit-innervating MNs. (H) Quantification of FOXP1^high^ and FOXP1^high^/SCIP MNs using FCS software. Staining intensity were extracted with Cell profiler and plotted using FCS. FOXP1 weak and negative cells were defined on the FGF D3-9 condition. The gate was then reported on the other conditions. (I) Proportion of FOXP1^high^ and FOXP1^high^/SCIP in upon modulation of FGF2 treatment duration. Scale bars: 100μm. Data are shown as mean ± SD.

**Figure S4.**
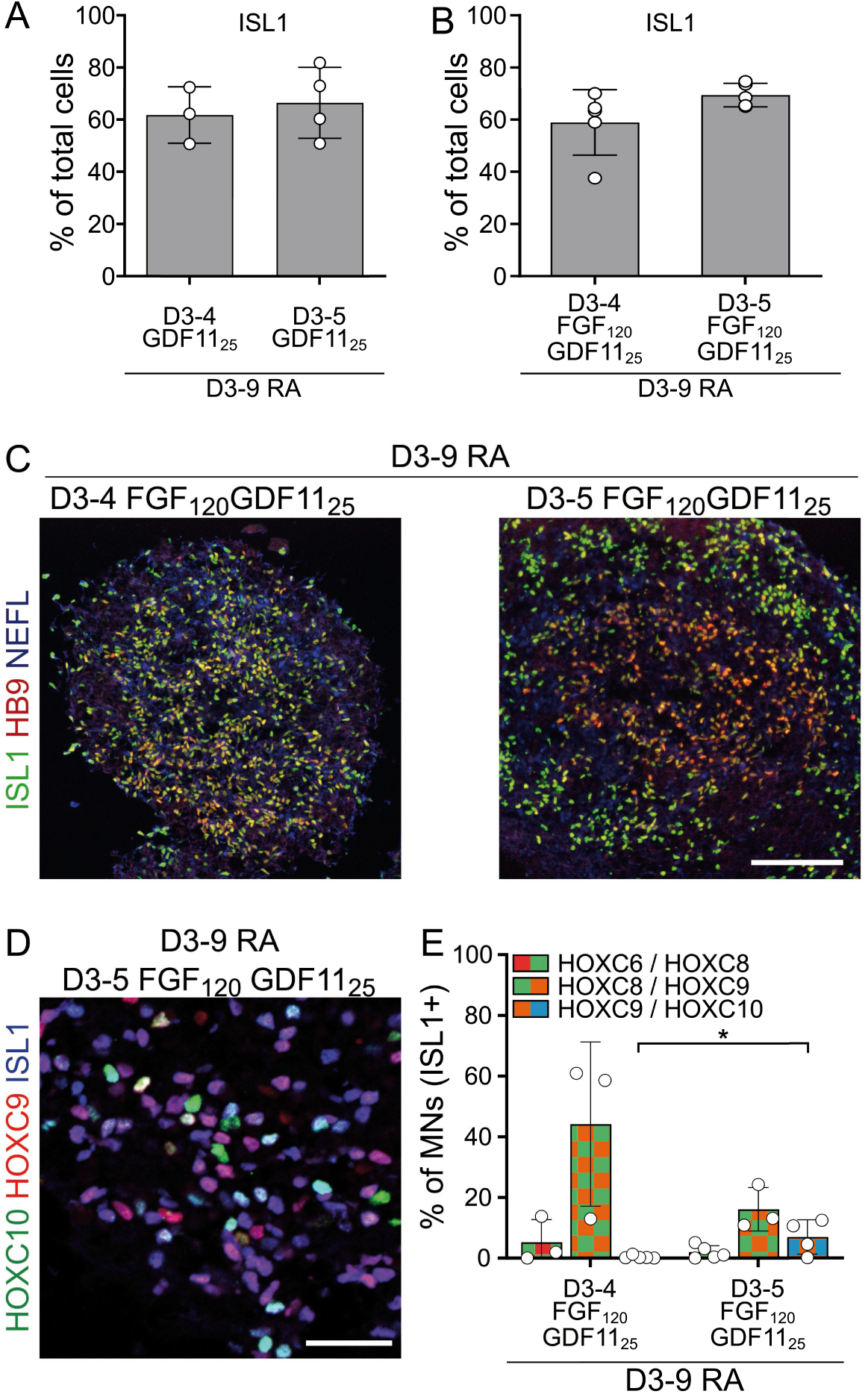
FGF2 and GDF11 cooperate to induce caudal thoracic and lumbar motor neurons. **(A)** Quantification of the percentage of MNs on day 14 of differentiation upon addition of GDF11. GDF11 (25 ng/ml) was added at day 3 for different durations. **(B)** Quantification of the percentage of MNs (ISL1+ cells) on day 14 of differentiation upon addition of FGF2 and GDF11. FGF2 (120 ng/ml) and GDF11 (25 ng/ml) were added on day 3 of differentiation for different durations. **(C)** Representative images of immunostainings for the motor neuron markers ISL1, HB9 and the pan neuronal marker NEFL on cryostat sections of EBs on day 14 of differentiation. Scale bar: 100μm **(D)** Representative images of immunostainings for ISL1, HOXC9 and HOXC10 on cryostat sections of EBs on day 14 of differentiation. Scale bar :30μm **(E)** Quantification of the percentage of HOXs co-expression in the indicated conditions.

**Table S1 - Transcriptomic data and list of markers defining axial progenitors and other developmentally related cell types**

The first panel displays the normalized number of reads for each gene for hESC, D2, D3 and D4 replicates. The following panels display the list of genes enriched (FC⩾2.0, p-value<0.05) in the indicated comparison. The last panel displays the list of genes that identify specific cell types (allantois, mesodermal progenitors) and the references used to construct these lists.

**Table S2.**
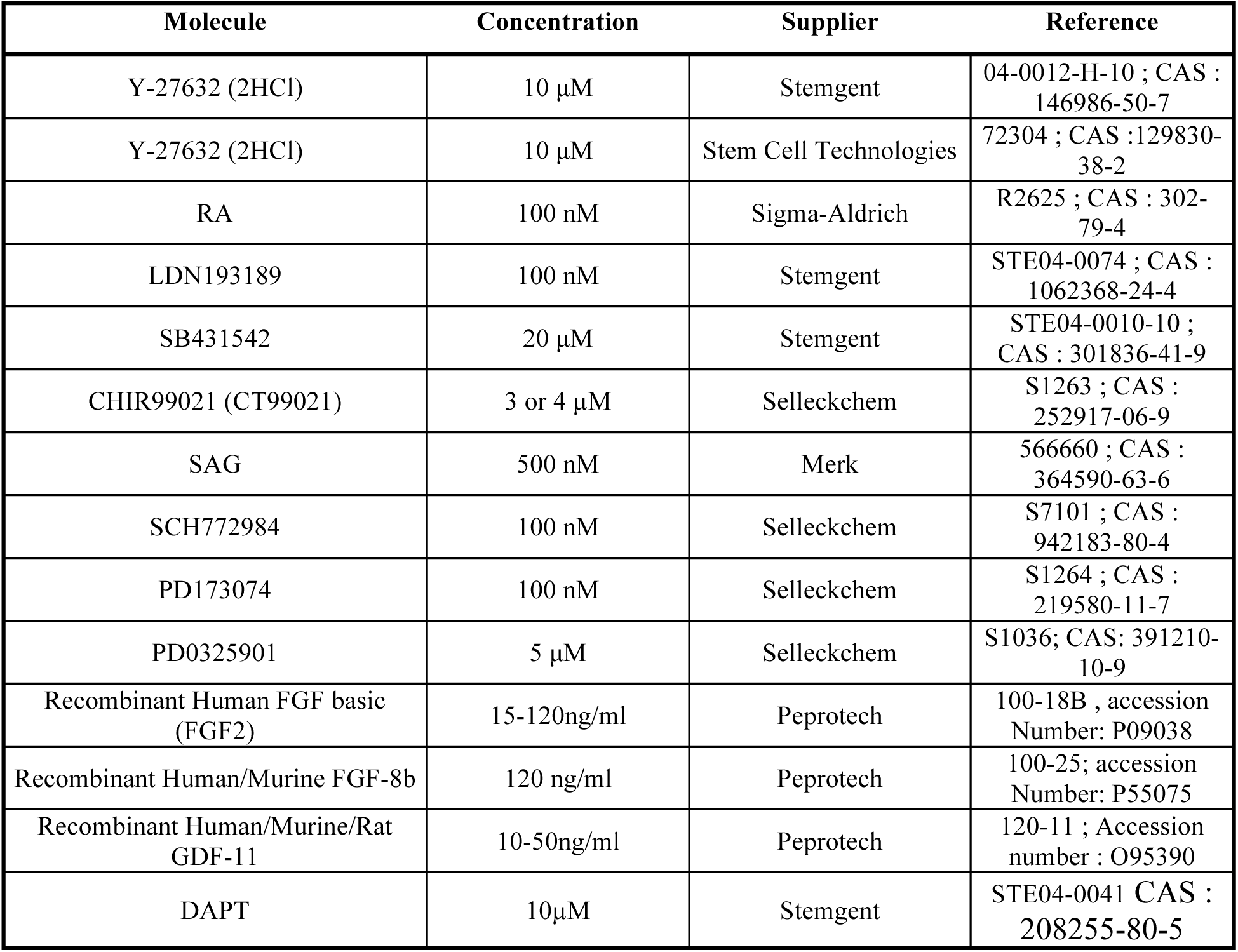
Growth factors and small molecules.

| Molecule | Concentration | Supplier | Reference |
| --- | --- | --- | --- |
| Y-27632 (2HCl) | 10 μM | Stemgent | 04-0012-H-10 ; CAS : 146986-50-7 |
| Y-27632 (2HCl) | 10 μM | Stem Cell Technologies | 72304 ; CAS : 129830-38-2 |
| RA | 100 nM | Sigma-Aldrich | R2625 ; CAS : 302-79-4 |
| LDN193189 | 100 nM | Stemgent | STE04-0074 ; CAS : 1062368-24-4 |
| SB431542 | 20 μM | Stemgent | STE04-0010-10 ; CAS : 301836-41-9 |
| CHIR99021 (CT99021) | 3 or 4 μM | Selleckchem | S1263 ; CAS : 252917-06-9 |
| SAG | 500 nM | Merk | 566660 ; CAS : 364590-63-6 |
| SCH772984 | 100 nM | Selleckchem | S7101 ; CAS : 942183-80-4 |
| PD173074 | 100 nM | Selleckchem | S1264 ; CAS : 219580-11-7 |
| PD0325901 | 5 μM | Selleckchem | S1036; CAS: 391210-10-9 |
| Recombinant Human FGF basic (FGF2) | 15-120ng/ml | Peprtech | 100-18B , accession Number: P09038 |
| Recombinant Human/Murine FGF-8b | 120 ng/ml | Peprtech | 100-25; accession Number: P55075 |
| Recombinant Human/Murine/Rat GDF-11 | 10-50ng/ml | Peprtech | 120-11 ; Accession number : O95390 |
| DAPT | 10μM | Stemgent | STE04-0041 CAS : 208255-80-5 |

**Table S3.** Antibodies.

| Target | HOST species | Dilution | References | Provider |
| --- | --- | --- | --- | --- |
| CDX2 | Rabbit | 1:1000 | AB76541<br>RRID: AB_1523334 | Abcam |
| FOXP1 | Rabbit | 1:32000 | AB16645<br>RRID: AB_732428 | Abcam |
| FOXP1 | Mouse | 1:3000 | MAB45341<br>RRID:AB_2107101 | R&D systems |
| HB9 | Mouse | 1:50 | 81.5C10<br>RRID: AB_2145209 | DSHB |
| HOXA5 | Rabbit | 1:1000 | ARP31447_P050<br>RRID:AB_2046241 | Sigma-Aldrich |
| HOXA5 | Guinea pig | 1 :1000 | Cu570 | H. Wichterle |
| HOXC5 | Rabbit | 1:1000 | HPA026794<br>RRID:AB_10612118 | Sigma-Aldrich |
| HOXC6 | Rabbit | 1:8000 | ARP38484_P50<br>RRID:AB_10866814 | Euromedex |
| HOXC8 | Mouse | 1:10000 | BLE920502 | Biologend |
| HOXC9 | Guinea pig | 1:300 | - | J. Dasen |
| HOXD9 | Rabbit | 1:1500 | SC-8320 | Santa-Cruz |
| HOXC10 | Rabbit | 1:800 | HPA053919<br>RRID: AB_2682308 | Sigma-Aldrich |
| HuC/D | Rabbit | 1:500 | A21271<br>RRID: AB_221448 | Invitrogen |
| ISL1 | Goat | 1:1000 | GT15051<br>RRID: AB_2126323 | Neuromics |
| NEUROFILAMENT | Chicken | 1:500 | CH22105<br>RRID: AB_2737102 | Neuromics |
| SCIP | Rabbit | 1:1000 | AB126746<br>RRID:AB_11130256 | Abcam |
| SOX2 | Mouse | 1:200 | AB79351<br>RRID:AB_10710406 | Abcam |
| TBXT/BRACHURY | Goat | 1:800 | AF2085<br>RRID:AB_2200235 | R&D systems |
| **Tested antibodies with no staining or unspecific one** |  |  |  |  |
| HOXC8 | Mouse |  | sc-517007 | Santa Cruz |
| HOXC9 | Rabbit |  | ARP35813 (Middle region)<br>RRID: AB_938161 | Aviva Sytems Biology |
| HOXC9 | Rabbit |  | H2041 n-terminal region<br>RRID: AB_10603946 | Sigma-Aldrich |
| HOXC9 | Rabbit |  | H1916<br>RRID: AB_10604162 | Sigma-Aldrich |
| HOXC10 | Mouse |  | 5G12-10 supernatant<br>RRID: AB_528286 | DSHB |

**Table S4.**
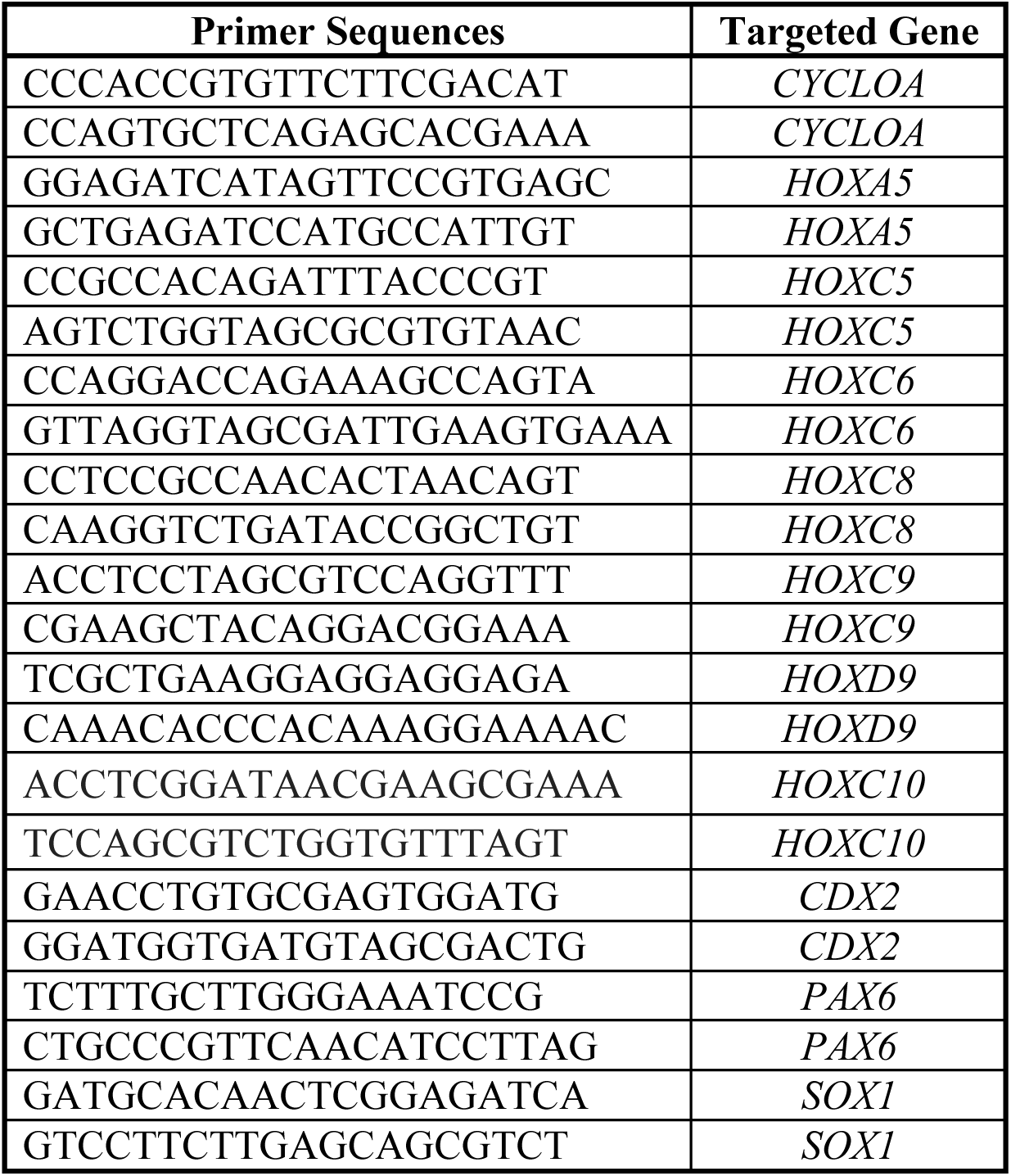
Real time PCR Primers.

## REFERENCES

Abati, E., Bresolin, N., Comi, G. and Corti, S. (2019). Advances, Challenges, and Perspectives in Translational Stem Cell Therapy for Amyotrophic Lateral Sclerosis. Mol. Neurobiol. 56, 6703–6715.

Aires, R., de Lemos, L., Nóvoa, A., Jurberg, A. D., Mascrez, B., Duboule, D. and Mallo, M. (2019). Tail Bud Progenitor Activity Relies on a Network Comprising Gdf11, Lin28, and Hox13 Genes. Dev. Cell 48, 383-395.e8.

Amoroso, M. W., Croft, G. F., Williams, D. J., O’Keeffe, S., Carrasco, M. A., Davis, A. R., Roybon, L., Oakley, D. H., Maniatis, T., Henderson, C. E., et al. (2013). Accelerated high-yield generation of limb-innervating motor neurons from human stem cells. J. Neurosci. 33, 574–586.

An, D., Fujiki, R., Iannitelli, D. E., Smerdon, J. W., Maity, S., Rose, M. F., Gelber, A., Wanaselja, E. K., Yagudayeva, I., Lee, J. Y., et al. (2019). Stem cell-derived cranial and spinal motor neurons reveal proteostatic differences between ALS resistant and sensitive motor neurons. Elife 8,.

Attardi, A., Fulton, T., Florescu, M., Shah, G., Muresan, L., Lenz, M. O., Lancaster, C., Huisken, J., van Oudenaarden, A. and Steventon, B. (2018). Neuromesodermal progenitors are a conserved source of spinal cord with divergent growth dynamics. Development 145, dev166728.

Bakooshli, M. A., Lippmann, E. S., Mulcahy, B., Iyer, N., Nguyen, C. T., Tung, K., Stewart, B. A., Van Den Dorpel, H., Fuehrmann, T., Shoichet, M., et al. (2019). A 3d culture model of innervated human skeletal muscle enables studies of the adult neuromuscular junction. Elife 8,.

Baloh, R. H., Glass, J. D. and Svendsen, C. N. (2018). Stem cell transplantation for amyotrophic lateral sclerosis. Curr. Opin. Neurol. 31, 655–661.

Bel-Vialar, S., Itasaki, N. and Krumlauf, R. (2002). Initiating Hox gene expression: In the early chick neural tube differential sensitivity to FGF and RA signaling subdivides the HoxB genes in two distinct groups. Development 129, 5103–5115.

Bell, S. W., Brown, M. J. C. and Hems, T. J. (2017). Refinement of myotome values in the upper limb: Evidence from brachial plexus injuries. Surgeon 15, 1–6.

Bialecka, M., Wilson, V. and Deschamps, J. (2010). Cdx mutant axial progenitor cells are rescued by grafting to a wild type environment. Dev. Biol. 347, 228–234.

Boulet, A. M. and Capecchi, M. R. (2012). Signaling by FGF4 and FGF8 is required for axial elongation of the mouse embryo. Dev. Biol. 371, 235–245.

Briscoe, J. and Novitch, B. G. (2008). Regulatory pathways linking progenitor patterning, cell fates and neurogenesis in the ventral neural tube. Philos. Trans. R. Soc. B Biol. Sci. 363, 57–70.

Cambray, N. and Wilson, V. (2007). Two distinct sources for a population of maturing axial progenitors. Development 134, 2829–2840.

Carpenter, A. E., Jones, T. R., Lamprecht, M. R., Clarke, C., Kang, I. H., Friman, O., Guertin, D. A., Chang, J. H., Lindquist, R. A., Moffat, J., et al. (2006). CellProfiler: Image analysis software for identifying and quantifying cell phenotypes. Genome Biol.

Catela, C., Shin, M. M., Lee, D. H., Liu, J. P. and Dasen, J. S. (2016). Hox Proteins Coordinate Motor Neuron Differentiation and Connectivity Programs through Ret/Gfrα Genes. Cell Rep. 14, 1901–1915.

Dasen, J. S. (2017). Master or servant? emerging roles for motor neuron subtypes in the construction and evolution of locomotor circuits. Curr. Opin. Neurobiol. 42, 25–32.

Dasen, J. S., Liu, J. P. and Jessell, T. M. (2003). Motor neuron columnar fate imposed by sequential phases of Hox-c activity. Nature 425, 926–933.

Dasen, J. S., De Camilli, A., Wang, B., Tucker, P. W. and Jessell, T. M. (2008). Hox Repertoires for Motor Neuron Diversity and Connectivity Gated by a Single Accessory Factor, FoxP1. Cell 134, 304–316.

Del Corral, R. D. and Storey, K. G. (2004). Opposing FGF and retinoid pathways: A signalling switch that controls differentiation and patterning onset in the extending vertebrate body axis. BioEssays 26, 857–869.

Denans, N., Iimura, T. and Pourquié, O. (2015). Hox genes control vertebrate body elongation by collinear Wnt repression. Elife 2015,.

Denham, M., Hasegawa, K., Menheniott, T., Rollo, B., Zhang, D., Hough, S., Alshawaf, A., Febbraro, F., Ighaniyan, S., Leung, J., et al. (2015). Multipotent caudal neural progenitors derived from human pluripotent stem cells that give rise to lineages of the central and peripheral nervous system. Stem Cells 33, 1759–1770.

Deschamps, J. and Duboule, D. (2017). Embryonic timing, axial stem cells, chromatin dynamics, and the Hox clock. Genes Dev. 31, 1406–1416.

Dias, J. M., Alekseenko, Z., Applequist, J. M. and Ericson, J. (2014). Tgfβ signaling regulates temporal neurogenesis and potency of neural stem cells in the CNS. Neuron 84, 927–939.

Diaz-Cuadros, M., Wagner, D. E., Budjan, C., Hubaud, A., Tarazona, O. A., Donelly, S., Michaut, A., Al Tanoury, Z., Yoshioka-Kobayashi, K., Niino, Y., et al. (2020). In vitro characterization of the human segmentation clock. Nature 580, 113–118.

Du, Z.-W., Chen, H., Liu, H., Lu, J., Qian, K., Huang, C.-L., Zhong, X., Fan, F. and Zhang, S.-C. (2015). Generation and expansion of highly pure motor neuron progenitors from human pluripotent stem cells. Nat. Commun. 6, 6626.

Durston, A. J. (2019). Some Questions and Answers About the Role of Hox Temporal Collinearity in Vertebrate Axial Patterning. Front. Cell Dev. Biol. 7,.

Duval, N., Vaslin, C., Barata, T. C., Frarma, Y., Contremoulins, V., Baudin, X., Nedelec, S. and Ribes, V. C. (2019). Bmp4 patterns smad activity and generates stereotyped cell fate organization in spinal organoids. Dev. 146, 24–33.

Ebisuya, M. and Briscoe, J. (2018). What does time mean in development? Development 145, dev164368.

Edri, S., Hayward, P., Jawaid, W. and Arias, A. M. (2019). Neuro-mesodermal progenitors (NMPs): A comparative study between pluripotent stem cells and embryo-derived populations. Dev. 146,.

Evtouchenko, L., Studer, L., Spencer, C., Dreher, E. and Seiler, R. W. (1996). A mathematical model for the estimation of human embryonic and fetal age. Cell Transplant. 5, 453–464.

Faustino Martins, J. M., Fischer, C., Urzi, A., Vidal, R., Kunz, S., Ruffault, P. L., Kabuss, L., Hube, I., Gazzerro, E., Birchmeier, C., et al. (2020). Self-Organizing 3D Human Trunk Neuromuscular Organoids. Cell Stem Cell 26, 172-186.e6.

Frith, T. J. R., Granata, I., Wind, M., Stout, E., Thompson, O., Neumann, K., Stavish, D., Heath, P. R., Ortmann, D., Hackland, J. O. S., et al. (2018). Human axial progenitors generate trunk neural crest cells in vitro. Elife 7,.

Gaunt, S. J. (1991). Expression patterns of mouse hox genes: Clues to an understanding of developmental and evolutionary strategies. BioEssays 13, 505–513.

Gaunt, S. J., George, M. and Paul, Y. L. (2013). Direct activation of a mouse Hoxd11 axial expression enhancer by Gdf11/Smad signalling. Dev. Biol. 383, 52–60.

Gouti, M., Tsakiridis, A., Wymeersch, F. J., Huang, Y., Kleinjung, J., Wilson, V. and Briscoe, J. (2014). In vitro generation of neuromesodermal progenitors reveals distinct roles for wnt signalling in the specification of spinal cord and paraxial mesoderm identity. PLoS Biol. 12, e1001937.

Gouti, M., Delile, J., Stamataki, D., Wymeersch, F. J., Huang, Y., Kleinjung, J., Wilson, V. and Briscoe, J. (2017). A Gene Regulatory Network Balances Neural and Mesoderm Specification during Vertebrate Trunk Development. Dev. Cell 41, 243-261.e7.

Henrique, D., Abranches, E., Verrier, L. and Storey, K. G. (2015). Neuromesodermal progenitors and the making of the spinal cord. Dev. 142, 2864–2875.

Izpisúa-Belmonte, J. C., Falkenstein, H., Dollé, P., Renucci, A. and Duboule, D. (1991). Murine genes related to the Drosophila AbdB homeotic genes are sequentially expressed during development of the posterior part of the body. EMBO J. 10, 2279–2289.

Jung, H., Lacombe, J., Mazzoni, E. O., Liem, K. F., Grinstein, J., Mahony, S., Mukhopadhyay, D., Gifford, D. K., Young, R. A., Anderson, K. V., et al. (2010). Global Control of Motor Neuron Topography Mediated by the Repressive Actions of a Single Hox Gene. Neuron 67, 781–796.

Jurberg, A. D., Aires, R., Varela-Lasheras, I., Nóvoa, A. and Mallo, M. (2013). Switching axial progenitors from producing trunk to tail tissues in vertebrate embryos. Dev. Cell 25, 451–462.

Kawaguchi, A. (2019). Temporal patterning of neocortical progenitor cells: How do they know the right time? Neurosci. Res. 138, 3–11.

Kimelman, D. and Martin, B. L. (2012). Anterior-posterior patterning in early development: Three strategies. Wiley Interdiscip. Rev. Dev. Biol. 1, 253–266.

Koch, F., Scholze, M., Wittler, L., Schifferl, D., Sudheer, S., Grote, P., Timmermann, B., Macura, K. and Herrmann, B. G. (2017). Antagonistic Activities of Sox2 and Brachyury Control the Fate Choice of Neuro-Mesodermal Progenitors. Dev. Cell 42, 514-526.e7.

Kohwi, M. and Doe, C. Q. (2013). Temporal fate specification and neural progenitor competence during development. Nat. Rev. Neurosci. 14, 823–838.

Lambrot, R., Coffigny, H., Pairault, C., Donnadieu, A. C., Frydman, R., Habert, R. and Rouiller-Fabre, V. (2006). Use of organ culture to study the human fetal testis development: Effect of retinoic acid. J. Clin. Endocrinol. Metab. 91, 2696–2703.

Li, X. J., Du, Z. W., Zarnowska, E. D., Pankratz, M., Hansen, L. O., Pearce, R. A. and Zhang, S. C. (2005). Specification of motoneurons from human embryonic stem cells. Nat. Biotechnol. 23, 215–221.

Lippmann, E. S., E.williams, C., Ruhl, D. A., Estevez-Silva, M. C., Chapman, E. R., Coon, J. J. and Ashton, R. S. (2015). Deterministic HOX patterning in human pluripotent stem cell-derived neuroectoderm. Stem Cell Reports 4, 632–644.

Liu, J. P. (2006). The function of growth/differentiation factor 11 (Gdf11) in rostrocaudal patterning of the developing spinal cord. Development 133, 2865–2874.

Liu, J. P., Laufer, E. and Jessell, T. M. (2001). Assigning the positional identity of spinal motor neurons: Rostrocaudal patterning of Hox-c expression by FGFs, Gdf11, and retinoids. Neuron 32, 997–1012.

Lunn, J. S., Fishwick, K. J., Halley, P. A. and Storey, K. G. (2007). A spatial and temporal map of FGF/Erk1/2 activity and response repertoires in the early chick embryo. Dev. Biol. 302, 536–552.

Machado, C. B., Kanning, K. C., Kreis, P., Stevenson, D., Crossley, M., Nowak, M., Iacovino, M., Kyba, M., Chambers, D., Blanc, E., et al. (2014). Reconstruction of phrenic neuron identity in embryonic stem cell-derived motor neurons. Dev. 141, 784–794.

Machado, C. B., Pluchon, P., Harley, P., Rigby, M., Sabater, V. G., Stevenson, D. C., Hynes, S., Lowe, A., Burrone, J., Viasnoff, V., et al. (2019). In Vitro Modeling of Nerve–Muscle Connectivity in a Compartmentalized Tissue Culture Device. Adv. Biosyst.

Mallo, M., Vinagre, T. and Carapuço, M. (2009). The road to the vertebral formula. Int. J. Dev. Biol. 53, 1469–1481.

Matsuda, M., Yamanaka, Y., Uemura, M., Osawa, M., Saito, M. K., Nagahashi, A., Nishio, M., Guo, L., Ikegawa, S., Sakurai, S., et al. (2020). Recapitulating the human segmentation clock with pluripotent stem cells. Nature 580, 124–129.

Maury, Y., Côme, J., Piskorowski, R. A., Salah-Mohellibi, N., Chevaleyre, V., Peschanski, M., Martinat, C. and Nedelec, S. (2015). Combinatorial analysis of developmental cues efficiently converts human pluripotent stem cells into multiple neuronal subtypes. Nat. Biotechnol. 33, 89–96.

Mazzoni, E. O., Mahony, S., Peljto, M., Patel, T., Thornton, S. R., McCuine, S., Reeder, C., Boyer, L. A., Young, R. A., Gifford, D. K., et al. (2013). Saltatory remodeling of Hox chromatin in response to rostrocaudal patterning signals. Nat. Neurosci. 16, 1191–1198.

McGrew, M. J., Sherman, A., Lillico, S. G., Ellard, F. M., Radcliffe, P. A., Gilhooley, H. J., Mitrophanous, K. A., Cambray, N., Wilson, V. and Sang, H. (2008). Localised axial progenitor cell populations in the avian tail bud are not committed to a posterior Hox identity. Development 135, 2289–2299.

McPherron, A. C., Lawler, A. M. and Lee, S. J. (1999). Regulation of anterior/posterior patterning of the axial skeleton by growth/differentiation factor 11. Nat. Genet. 22, 260–264.

Mendelsohn, A. I., Dasen, J. S. and Jessell, T. M. (2017). Divergent Hox Coding and Evasion of Retinoid Signaling Specifies Motor Neurons Innervating Digit Muscles. Neuron 93, 792-805.e4.

Metzis, V., Steinhauser, S., Pakanavicius, E., Gouti, M., Stamataki, D., Ivanovitch, K., Watson, T., Rayon, T., Mousavy Gharavy, S. N., Lovell-Badge, R., et al. (2018). Nervous System Regionalization Entails Axial Allocation before Neural Differentiation. Cell 175, 1105-1118.e17.

Narendra, V., Rocha, P. P., An, D., Raviram, R., Skok, J. A., Mazzoni, E. O. and Reinberg, D. (2015). CTCF establishes discrete functional chromatin domains at the Hox clusters during differentiation. Science (80-.). 347, 1017–1021.

Neijts, R., Amin, S., van Rooijen, C. and Deschamps, J. (2017). Cdx is crucial for the timing mechanism driving colinear Hox activation and defines a trunk segment in the Hox cluster topology. Dev. Biol. 422, 146–154.

Nijssen, J., Comley, L. H. and Hedlund, E. (2017). Motor neuron vulnerability and resistance in amyotrophic lateral sclerosis. Acta Neuropathol. 133, 863–885.

Noordermeer, D., Leleu, M., Splinter, E., Rougemont, J., De Laat, W. and Duboule, D. (2011). The dynamic architecture of Hox gene clusters. Science (80-.). 334, 222–225.

Noordermeer, D., Leleu, M., Schorderet, P., Joye, E., Chabaud, F. and Duboule, D. (2014). Temporal dynamics and developmental memory of 3D chromatin architecture at Hox gene loci. Elife 2014,.

Oberst, P., Fièvre, S., Baumann, N., Concetti, C., Bartolini, G. and Jabaudon, D. (2019a). Temporal plasticity of apical progenitors in the developing mouse neocortex. Nature 573, 370–374.

Oberst, P., Agirman, G. and Jabaudon, D. (2019b). Principles of progenitor temporal patterning in the developing invertebrate and vertebrate nervous system. Curr. Opin. Neurobiol. 56, 185–193.

Ogura, T., Sakaguchi, H., Miyamoto, S. and Takahashi, J. (2018). Three-dimensional induction of dorsal, intermediate and ventral spinal cord tissues from human pluripotent stem cells. Development 145, dev162214.

Olivera-Martinez, I., Harada, H., Halley, P. A. and Storey, K. G. (2012). Loss of FGF-Dependent Mesoderm Identity and Rise of Endogenous Retinoid Signalling Determine Cessation of Body Axis Elongation. PLoS Biol. 10,.

Osaki, T., Uzel, S. G. M. and Kamm, R. D. (2020). On-chip 3D neuromuscular model for drug screening and precision medicine in neuromuscular disease. Nat. Protoc. 15, 421–449.

Peljto, M., Dasen, J. S., Mazzoni, E. O., Jessell, T. M. and Wichterle, H. (2010). Functional diversity of ESC-derived motor neuron subtypes revealed through intraspinal transplantation. Cell Stem Cell 7, 355–366.

Philippidou, P. and Dasen, J. S. (2013). Hox Genes: Choreographers in Neural Development, Architects of Circuit Organization. Neuron 80, 12–34.

Philippidou, P., Walsh, C. M., Aubin, J., Jeannotte, L. and Dasen, J. S. (2012). Sustained Hox5 gene activity is required for respiratory motor neuron development. Nat. Neurosci. 15, 1636–1644.

Pourquié, O., Al Tanoury, Z. and Chal, J. (2018). The Long Road to Making Muscle In Vitro. In Current Topics in Developmental Biology, pp. 123–142.

Ragagnin, A. M. G., Shadfar, S., Vidal, M., Jamali, M. S. and Atkin, J. D. (2019). Motor neuron susceptibility in ALS/FTD. Front. Neurosci. 13,.

Ribes, V., Le Roux, I., Rhinn, M., Schuhbaur, B. and Dolle, P. (2009). Early mouse caudal development relies on crosstalk between retinoic acid, Shh and Fgf signalling pathways. Development 136, 665–676.

Rossi, A. M., Fernandes, V. M. and Desplan, C. (2017). Timing temporal transitions during brain development. Curr. Opin. Neurobiol. 42, 84–92.

Rousso, D. L., Gaber, Z. B., Wellik, D., Morrisey, E. E. and Novitch, B. G. (2008). Coordinated Actions of the Forkhead Protein Foxp1 and Hox Proteins in the Columnar Organization of Spinal Motor Neurons. Neuron 59, 226–240.

Routal, R. V. and Pal, G. P. (1999). A study of motoneuron groups and motor columns of the human spinal cord. J. Anat. 195, 211–224.

Sances, S., Bruijn, L. I., Chandran, S., Eggan, K., Ho, R., Klim, J. R., Livesey, M. R., Lowry, E., Macklis, J. D., Rushton, D., et al. (2016). Modeling ALS with motor neurons derived from human induced pluripotent stem cells. Nat. Neurosci. 19, 542–553.

Schindelin, J., Arganda-Carreras, I., Frise, E., Kaynig, V., Longair, M., Pietzsch, T., Preibisch, S., Rueden, C., Saalfeld, S., Schmid, B., et al. (2012). Fiji: An open-source platform for biological-image analysis. Nat. Methods 9, 676–682.

Soshnikova, N. and Duboule, D. (2009). Epigenetic temporal control of mouse hox genes in vivo. Science (80-.). 324, 1321–1323.

Steinbeck, J. A. and Studer, L. (2015). Moving stem cells to the clinic: Potential and limitations for brain repair. Neuron 86, 187–206.

Steinbeck, J. A., Choi, S. J., Mrejeru, A., Ganat, Y., Deisseroth, K., Sulzer, D., Mosharov, E. V. and Studer, L. (2015). Optogenetics enables functional analysis of human embryonic stem cell-derived grafts in a Parkinson’s disease model. Nat. Biotechnol. 33, 204–209.

Syed, M. H., Mark, B. and Doe, C. Q. (2017). Playing Well with Others: Extrinsic Cues Regulate Neural Progenitor Temporal Identity to Generate Neuronal Diversity. Trends Genet. 33, 933–942.

Tiberi, L., Vanderhaeghen, P. and van den Ameele, J. (2012). Cortical neurogenesis and morphogens: Diversity of cues, sources and functions. Curr. Opin. Cell Biol. 24, 269–276.

Tschopp, P., Tarchini, B., Spitz, F., Zakany, J. and Duboule, D. (2009). Uncoupling time and space in the collinear regulation of Hox genes. PLoS Genet. 5,.

Tung, Y. T., Peng, K. C., Chen, Y. C., Yen, Y. P., Chang, M., Thams, S. and Chen, J. A. (2019). Mir-17∼92 Confers Motor Neuron Subtype Differential Resistance to ALS-Associated Degeneration. Cell Stem Cell 25, 193-209.e7.

Tzouanacou, E., Wegener, A., Wymeersch, F. J., Wilson, V. and Nicolas, J. F. (2009). Redefining the Progression of Lineage Segregations during Mammalian Embryogenesis by Clonal Analysis. Dev. Cell 17, 365–376.

Verrier, L., Davidson, L., Gierlinski, M., Dady, A. and Storey, K. G. (2018). Neural differentiation, selection and transcriptomic profiling of human neuromesodermal progenitor-like cells in vitro. Dev. 145, dev166215.

Wang, J., Vasaikar, S., Shi, Z., Greer, M. and Zhang, B. (2017). WebGestalt 2017: A more comprehensive, powerful, flexible and interactive gene set enrichment analysis toolkit. Nucleic Acids Res. 45, W130–W137.

Wymeersch, F. J., Huang, Y., Blin, G., Cambray, N., Wilkie, R., Wong, F. C. K. and Wilson, V. (2016). Position-dependent plasticity of distinct progenitor types in the primitive streak. Elife 5,.

Wymeersch, F. J., Skylaki, S., Huang, Y., Watson, J. A., Economou, C., Marek-Johnston, C., Tomlinson, S. R. and Wilson, V. (2019). Transcriptionally dynamic progenitor populations organised around a stable niche drive axial patterning. Dev. 146, dev168161.

Young, T., Rowland, J. E., van de Ven, C., Bialecka, M., Novoa, A., Carapuco, M., van Nes, J., de Graaff, W., Duluc, I., Freund, J. N., et al. (2009). Cdx and Hox Genes Differentially Regulate Posterior Axial Growth in Mammalian Embryos. Dev. Cell.

